# Structural basis for Potassium transport by KdpFABC

**DOI:** 10.1101/2021.01.09.426067

**Authors:** Marie E. Sweet, Casper Larsen, Xihui Zhang, Michael Schlame, Bjørn P. Pedersen, David L. Stokes

## Abstract

KdpFABC is an oligomeric K^+^ transport complex in prokaryotes that maintains ionic homeostasis under stress conditions. The complex comprises a channel-like subunit (KdpA) from the Superfamily of K^+^ Transporters and a pump-like subunit (KdpB) from the superfamily of P-type ATPases. Recent structural work has defined the architecture and generated contradictory hypotheses for the transport mechanism. Here, we use substrate analogs to stabilize four key intermediates in the reaction cycle and determine the corresponding structures by cryo-EM. We find that KdpB undergoes conformational changes consistent with other representatives from the P-type superfamily, whereas KdpA, KdpC and KdpF remain static. We observe a series of spherical densities that we assign as K^+^ or water and which define a pathway for K^+^ transport. This pathway runs through an intramembrane tunnel in KdpA and delivers ions to sites in the membrane domain of KdpB. Our structures suggest a mechanism where ATP hydrolysis is coupled to K^+^ transfer between alternative sites in the membrane domain of KdpB, ultimately reaching a low-affinity site where a water-filled pathway allows release of K^+^ to the cytoplasm.

## Introduction

KdpFABC is an ATP-dependent K^+^ pump in prokaryotes, essential for osmoregulation in K^+^-deficient environments. Expression of kdpFABC is induced when external K^+^ concentrations fall into the micromolar range, where constitutive K^+^-uptake systems, Trk and Kup, can no longer maintain intracellular K^+^ levels. Under these conditions, a high-affinity active transport system is required to maintain internal concentrations of K^+^, essential for regulating pH, membrane potential and the turgor pressure that drives cell growth and division (Altendorf et al., 2009). As a molecular machine, the oligomeric KdpFABC complex represents a fascinating hybrid that couples a channel-like subunit (KdpA) - related to the Superfamily of K^+^ Transporters (SKT) - with a pump-like subunit (KdpB) - belonging to the superfamily of P-type ATPases (Greie, 2011). Early studies established a role for KdpA in the selectivity and transport of K^+^ (Buurman et al., 1995). Furthermore, analysis of KdpA sequence and topology established the existence of four approximate repeats of the “MPM” fold that characterizes K^+^ channels, and a close resemblance to bacterial channels TrkH and KtrB (Durell et al., 2000). KdpB, on the other hand, harnesses the energy of ATP to drive the transport process. KdpB was shown to employ the Post-Albers reaction scheme (Siebers and Altendorf, 1989) that features two main conformational states, E1 and E2, and an aspartyl phosphate intermediate (Fig. 1b). However, mechanisms for energy coupling between these two subunits have remained elusive, as has the specific transport pathway of K^+^ through the complex (Pedersen et al., 2019).

**Fig 1.**
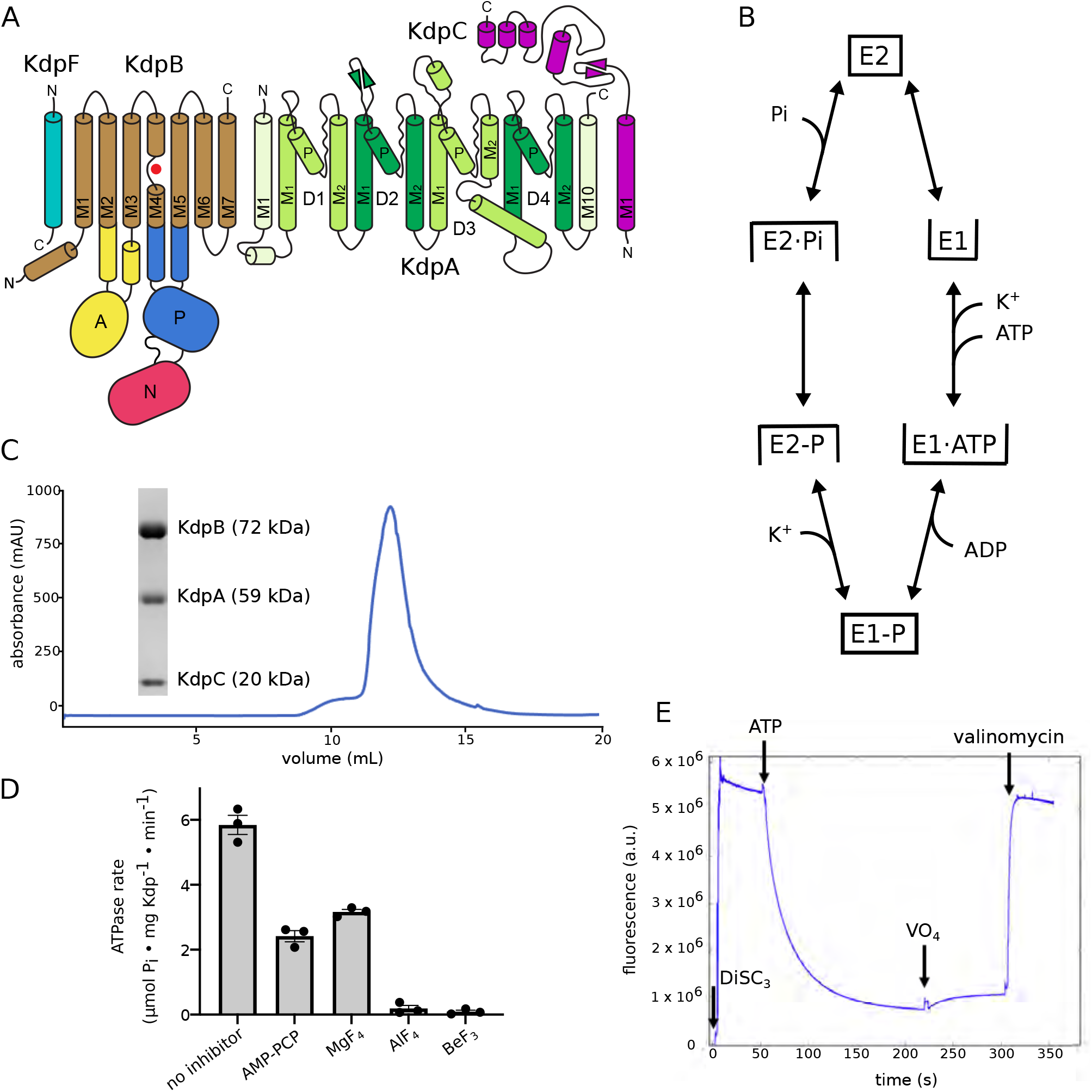
Topology, purification and activity of KdpFABC. **(A)** Topology of KdpFABC complex. KdpA (green) is characterized by four pseudorepeats (D1-D4) of the MPM motif. The selectivity filter is formed by the loops between the pore helix (P) helix and M2. KdpB has seven membrane helices (brown) with a water bound to M4 and three cytoplasmic domains denoted A (yellow), N (red) and P (blue). KdpC (purple) and KdpF (cyan) have a single membrane helix. **(B)** Schematic of the Post-Albers reaction cycle used by P-type ATPases. Closed boxes indicate occluded states, whereas upward facing boxes are open to the periplasm and downward facing boxes are open to the cytoplasm. **(C)** Purity of KdpFABC preparations used for this work is illustrated by SDS-PAGE and a size-exclusion chromatogram. KdpF is not visible on the gel due to its small size. **(D)** Effect of substrate analogs on ATPase activity of KdpFABC showing partial inhibition from 5 mM AMP-PCP and 1 mM MgF_4_, but complete inhibition by 1 mM AlF_4_, and 2.5 mM BeF_3_ under turnover conditions. **(E)** Transport assay using the voltage-sensitive DiSC_3_ dye demonstrates robust activity of the construct used for this work. Addition of ATP leads to rapid buildup of membrane potential, which is collapsed upon addition of the K^+^ ionophore, valinomycin. Partial inhibition by VO_4_ is also reflected in a shift in membrane potential.

Recent structural studies have generated renewed interest in KdpFABC and have led to new and conflicting ideas about its transport process. The first structure was determined by X-ray crystallography, defining the architecture of the complex (Huang et al., 2017). This structure revealed a water molecule bound at the unwound part of the M4 transmembrane helix of KdpB (Fig. 1a). This site, here denoted Bx, is conserved among P-type ATPases and plays a key role in binding and transporting their respective substrates. Structures of SERCA revealed ions at two sites, dubbed site Ca(I) and site Ca(II), the latter of which is congruent with the Bx site in KdpB. Structures of Na,K-ATPase show a similar ion binding pattern, with a third Na^+^ ion located somewhat displaced from these two conserved sites. In KdpA, a K^+^ ion was bound within the selectivity filter (SF) and, like TrkH and KtrB, the third MPM repeat featured a kinked helix (D3M_2_ in Fig. 1a) with a loop that blocks this ion from traveling across the membrane. This loop was suggested to function as a gate that would move aside to open the pore for transport. In addition, a 40 Å long tunnel was seen encapsulated within the membrane domain of KdpFABC. This intramembrane tunnel connects the SF in KdpA with the Bx site in KdpB, leading to a novel proposal for energy coupling based on a Grothuss mechanism for charge transfer between these sites. According to this mechanism, the tunnel is filled with water molecules acting as a water wire to shuttle protons between the subunits. K^+^ binding to the SF would initiate proton hopping through the tunnel. Arrival of this charge at the Bx site would stimulate the cycle of ATP hydrolysis, as documented in other P-type ATPases. A salt-bridge network connecting cytoplasmic loops of KdpA and KdpB would then tug on the kinked D3M_2_ helix in KdpA, thus displacing the gating loop and allowing K^+^ to transit the membrane.

Two additional structures of KdpFABC were subsequently determined by cryo-EM, revealing two new conformations (Stock et al., 2018). Whereas the crystal structure appeared to represent an inhibited conformation, these cryo-EM structures were compatible with E1 and E2 states. An intact intramembrane tunnel was observed in the E1 state, but was interrupted in the E2 state due to conformational changes in KdpB. These observations led to proposal of an alternative mechanism in which K^+^ ions move through the tunnel, from the SF of KdpA to the Bx site of KdpB, where they are released to the cytoplasm at the appropriate step in the reaction cycle. The other two subunits, KdpC and KdpF, were also observed in all of these structures. KdpF consists of a single transmembrane helix at the interface of KdpA and KdpB, possibly serving to stabilize the complex. KdpC has a unique periplasmic domain anchored by a single TM helix; the location at the entrance to the SF suggested that this domain might act as a periplasmic filter or gate, though evidence for this role is currently lacking.

To shed more light on the coupling between KdpA and KdpB and to resolve the role of the intramembrane tunnel, we determined structures of the KdpFABC complex in all of the major enzymatic states. We used inhibitors to trap the complex in various discrete states, both in the presence and absence of K^+^, and imaged these samples by cryo-EM. In this way, we produced 14 independent density maps at resolutions between 2.9 and 3.7 Å; four of these were selected to represent the primary intermediates from the reaction cycle. These resulting structures display significant conformational changes in KdpB as well as non-protein spherical densities within the SF of KdpA, within the intramembrane tunnel, and in KdpB near the Bx site. By providing high-resolution detail to structural changes in the membrane domain of KdpB during the reaction cycle, we provide evidence and a comprehensive model for the transport mechanism in which K^+^ from the periplasm enters the SF of KdpA, moves through the tunnel to the Bx site, and is released to the cytoplasm by KdpB.

## Results

### Cryo-EM structures reveal conformational changes of KdpFABC

For this work, we introduced the Q116R mutation into KdpA and the S162A mutation into KdpB. As previously reported (Sweet et al., 2020), the resulting protein is fully functional and offers practical advantages for comparative study of reaction intermediates. Whereas the wild-type protein features very high (micromolar) affinity for K^+^, the Q116R mutation at the mouth of the SF lowers this affinity by three orders of magnitude, and has been used historically for comparison between K^+^-bound and K^+^-free states (Epstein et al., 1978; Siebers and Altendorf, 1989). The S162A mutation prevents phosphorylation of this serine, recently shown to be a regulatory modification that inhibits KdpFABC (Sweet et al., 2020). We expressed this mutant complex in *E. coli* and purified it in n-decyl-β-D-maltoside detergent using affinity chromatography followed by size-exclusion chromatography (Fig. 1c). The activity of the purified protein complex was documented both with ATPase assays of the detergent-solubilized samples and with K^+^ transport assays after reconstitution into lipid vesicles (Fig. 1d,e). For cryo-EM imaging, aliquots of the detergent-solubilized samples were mixed with various substrate analogs followed by rapid freezing of EM grids. Work on other P-type ATPases has identified a variety of substrate analogs that trap specific reaction intermediates (Møller et al., 2010). Accordingly, AMP-PCP was used to stabilize the E1 state that precedes formation of the covalent aspartyl phosphate (E1∼P), and four phosphate analogs were used to trap different E2 sub-states. BeF_3_ has been shown previously to trap the pre-hydrolysis E2-P state, AlF_4_ the transition state E2∼P, and MgF_4_ the post-hydrolysis E2·Pi state in other P-type ATPases (Danko and Suzuki, 2016). Orthovanadate is another phosphate analog with a long history of inhibiting P-type ATPases (Pedersen and Carafoli, 1987). ATPase assays confirmed the inhibitory capabilities of these various compounds (Fig. 1d), though it should be noted that turnover conditions with the presence of ATP and its reaction products are not truly representative of the inhibitory complexes produced for cryo-EM. For all but one substrate analog (MgF_4_), we prepared samples both in the presence and absence of 100 mM K^+^.

A total of 14 independent density maps were produced from the various samples using conventional methods of single-particle cryo-EM. Each data set comprised between 150,000 - 450,000 particles and overall resolutions ranged between 2.9 Å and 3.7 Å. In two cases, different conformations were resolved from single datasets using 3D classification. In the first case, samples treated with VO_4_ produced structures of both E1 and E2 states, suggesting that this phosphate analog has relatively low affinity and binds to only a fraction of the molecules. In the second case, treatment with BeF_3_ in the presence of K^+^ also produced two distinct maps, but these appear to be pre- and post-hydrolysis E2-P states, as is discussed in the next section. From the 14 density maps, we chose the highest resolution examples of the four unique conformational states that were observed, which were then used for model building and further detailed analysis. In addition, a second structure for the E2·Pi state was built for a direct comparison of structures in the presence and absence of K^+^ (Suppl. Figs 1-4, Suppl. Table 1).

Density maps representing the four main states in the reaction cycle are shown in Fig. 2 together with close-up views of the catalytic site of KdpB, where the substrate analogs bind. In the absence of substrates at the catalytic site (apo state), the A- and N-domains of KdpB are quite disordered, reflecting the innate mobility of these domains, which is explicitly depicted by an analysis of variability in the dataset (Suppl. Movies 1-3). Binding of ATP or phosphate analogs to the catalytic site stabilizes interactions between these domains and the P-domain, which carries the catalytic aspartate (Asp307). Densities for the analogs are clearly visible together with the key coordinating residues, except for those in the E2-P map which had lower resolution in this region. Overall, KdpA, KdpC, KdpF and the membrane domain of KdpB are well ordered, with clear visibility of side chain density.

**Fig 2.**
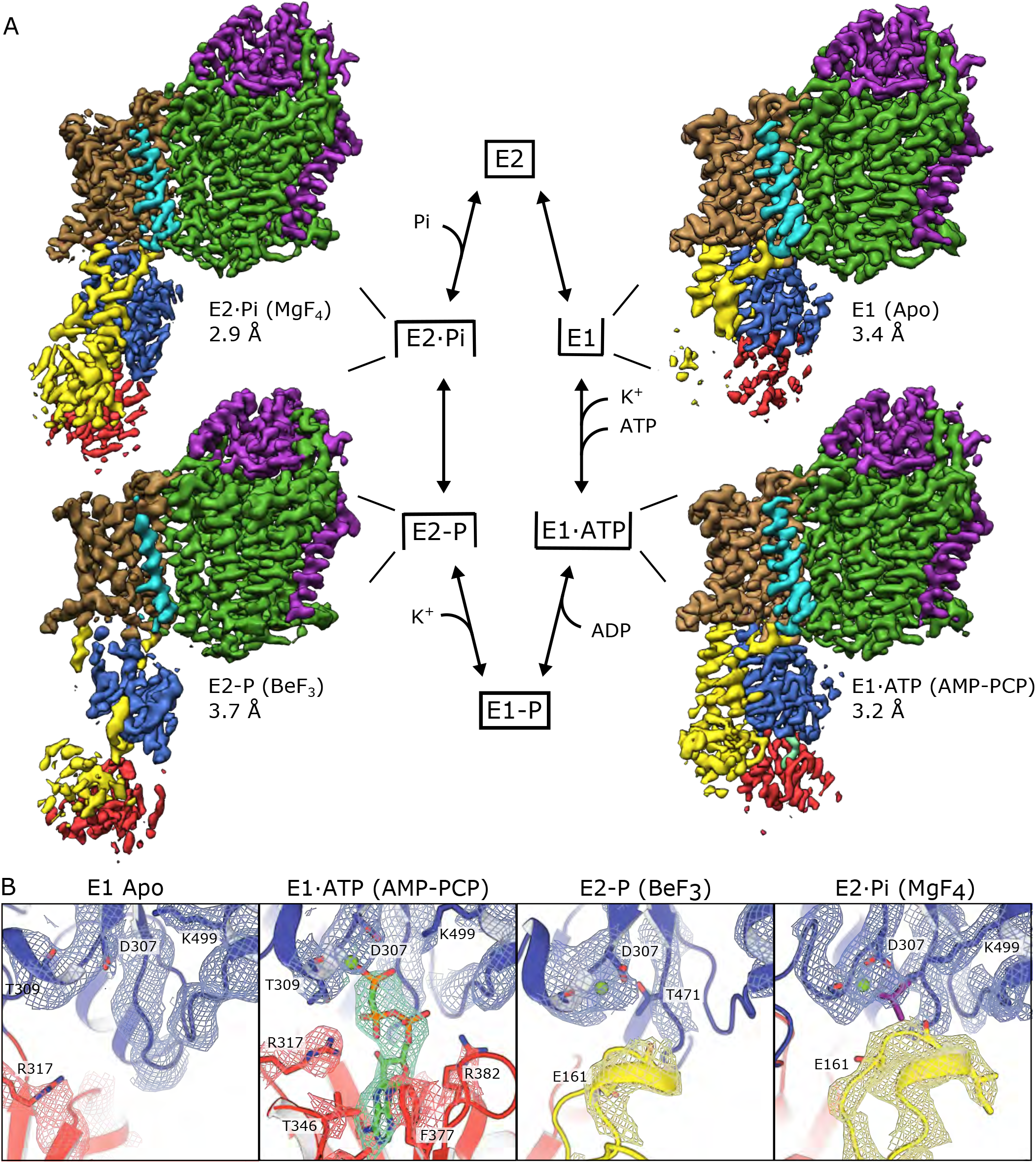
Overview of cryo-EM structures. **(A)** Cryo-EM density maps of KdpFABC stabilized in four unique conformational states, corresponding to intermediates in the Post-Albers reaction cycle. The complex is colored according to the scheme in Fig. 1. Density for AMP-PCP bound in the E1·ATP state is shown in cyan. **(B)** Closeup of catalytic sites with atomic models built for the density maps (mesh) shown in A. Density is clearly visible for AMP-PCP and MgF_4_; lower resolution of the E2-P map makes explicit placement of BeF_3_ ambiguous. Mg^2+^ ions shown in green and domains colored as in A.

### Conformations of KdpB confirm the stabilization of specific reaction intermediates

The reaction intermediate represented by each structure was confirmed by comparing KdpB with the well-studied P-type ATPase SERCA. After alignment of P-domains, N- and A-domain placements are consistent with specific states of SERCA: E1 state (PDBID: 1SU4), E1·ATP state (PDBID: 3N8G), E2-P state (PDBID: 3B9B), and E2·Pi state (PDBID: 2O9J) (Suppl. Fig. 5a). The E1 states show unliganded catalytic aspartic acids (Asp307 in KdpB and Asp351 in SERCA) that do not interact with either A- or N-domains, allowing them to retain their innate flexibility. The E1·ATP states show AMP-PCP molecules bound in equivalent pockets of the N-domains with the γ-phosphate poised for phosphoryl-transfer; conserved residues coordinate the Mg^2+^ and the ATP analog in an analogous fashion, and the A-domain is displaced from the catalytic site (Suppl. Fig. 5b). It is notable that the A-domain adopts a different angle in the E1·ATP state: the conserved TGES loop is rotated clockwise from the catalytic site in KdpB, whereas this loop is rotated counter-clockwise in SERCA as viewed from the cytoplasm. The pre-hydrolysis (E2-P) and post-hydrolysis (E2·Pi) E2 states in both SERCA and KdpB have a rather similar configurations at the catalytic site with the conserved TGES motif engaging the catalytic aspartate and displacing the N-domain. Although these two E2 states are difficult to distinguish solely on catalytic site residues, comparison of the linkage between cytoplasmic and membrane domains proves decisive. Specifically, the cytoplasmic extension of M2 becomes unwound in the transition between the E2-P and E2·Pi state of SERCA, producing a large scale structural difference that affects the position of cytoplasmic domains relative the membrane domain (Suppl. Fig. 5c). KdpB undergoes an analogous change in which M2 is continuous through Lys98 in E2-P, but becomes unwound after Leu85 in E2·Pi. As a result, the cytoplasmic domains move in a similar way and the TGES loop is repositioned to reflect its role in hydrolysis of the aspartyl phosphate. These comparisons confirm that similar chemistry is at play in the reaction cycles of these two homologs, and that substrate analogs stabilize KdpFABC in comparable enzymatic states from the Post-Albers cycle.

### KdpA, KdpC, and KdpF and structural effects of K^+^

In contrast to the large-scale conformational changes in KdpB, the other subunits of the complex are essentially unchanged as the molecule transitions between the different enzymatic states. Pairwise alignment of KdpA and KdpC produces RMSD’s <0.5 Å for Cα atoms, which reflects virtually no changes in the core structure and only slight changes in the peripheral loops of these subunits (Fig. 3a). These subunits are also unchanged compared to the crystal structure (RMSD’s < 1.0), indicating that they remain rigid throughout the reaction cycle. In the E1 states, a salt-bridge network exists between cytoplasmic loops on KdpA (Arg400 and Ser517) and KdpB (Asp300 and Asp302, Suppl. Fig. 6a). According to our previous mechanistic model based on the crystal structure, these interactions play a role in energy coupling by pulling on a coupling helix (D3M_2_ colored pink in Suppl. Fig. 6a) and opening a gate below the SF in KdpA (Huang et al., 2017). In order to test this model, we introduced the D300A/D302A mutation into the wild-type gene for kdpB and evaluated ATPase and transport activities after reconstitution of the purified complex into liposomes (Suppl. Fig. 6). The results show that both constructs remain coupled, with robust transport activity, although the mutations reduce V_max_ by 20-30%. For the transport assay, we also evaluated ATP dependence, thus deducing K_M_ of 5.7 and 13.9 μM for wild-type and mutant protein, respectively. Given the modest effects of the D300A/D302A mutation, we conclude that this salt-bridge network is not likely to be crucial for the transport mechanism, suggesting that an alternative transport mechanism is likely.

**Fig 3.**
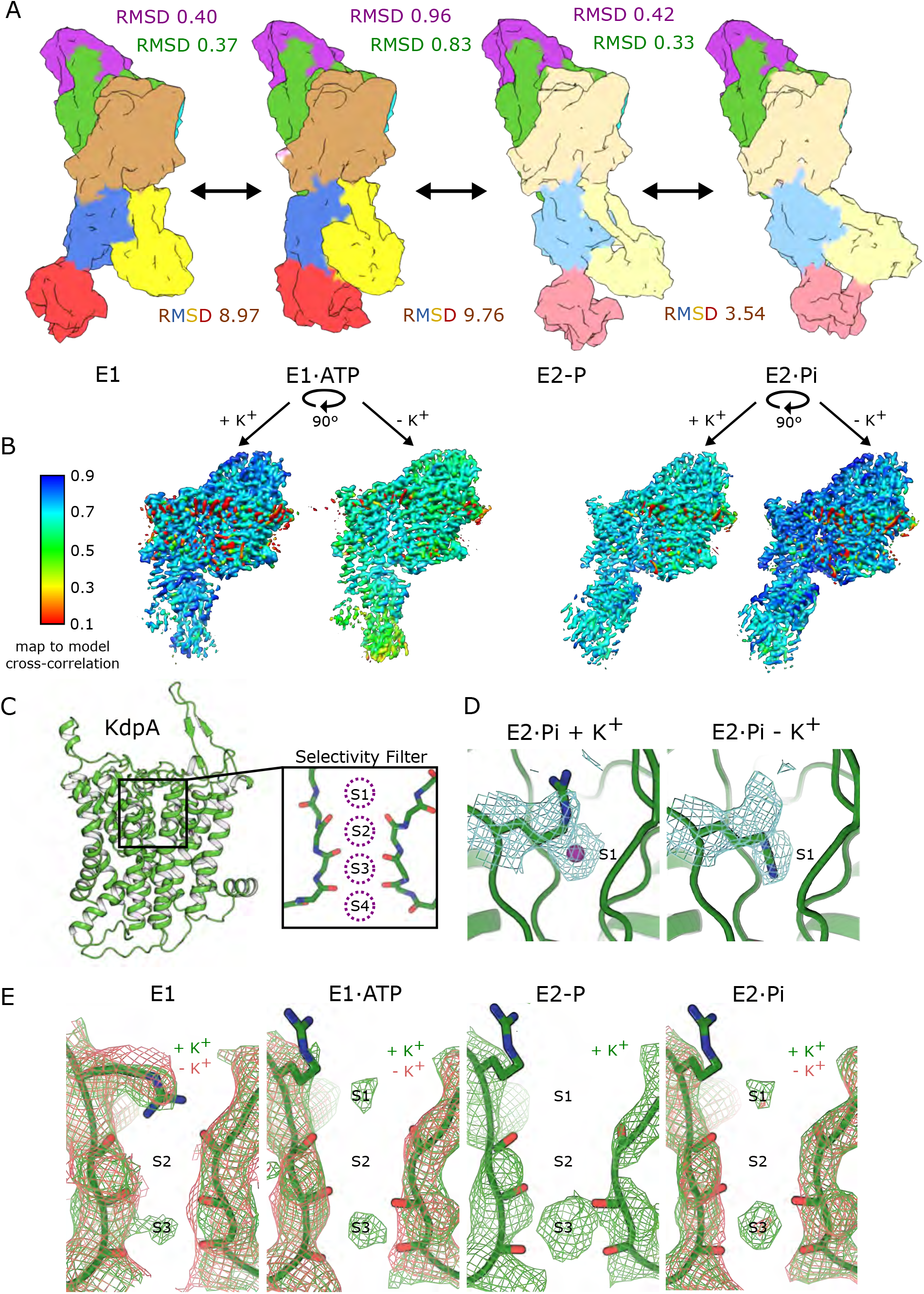
Conformational changes accompany the reaction cycle. **(A)** Surface representations of the KdpFABC complex rendered at 10 Å resolution show large scale conformational changes associated with the different states. Subunits are colored as in previous figures, though colors are more saturated in the E1 states to distinguish them from E2 states. RMSDs reflect comparison between individual subunits from adjacent conformational states (557, 672, 186 Cα atoms for KdpA, KdpB, KdpC, respectively). Text color indicates the subunit being compared (purple for KdpC, green for KdpA, brown/blue/yellow/red for KdpB). These comparisons indicate very little change in KdpA and KdpC, but large-scale changes in KdpB. **(B)** Comparison of E1·ATP and E2·Pi states in the presence and absence of K^+^ illustrates little to no effect of this ion on the conformation. Colors indicate model-to-map cross-correlation: in each case, the atomic model was built for the map produced in the presence of K^+^. Small red densities near the membrane domain arise from the surrounding micelle. **(C)** Cartoon representation of KdpA in the E1 state with an inset depicting the SF and four binding sites for K^+^ (S1-S4) observed in canonical K^+^ channels. **(D)** S1 site in the SF with map densities (contoured at 12σ) associated with Arg116 in the E2·Pi state in the presence and absence of K^+^. The bifurcated density suggests that this side chain competes with K^+^ at this site. **(E)** Map densities within the SF in the presence (green mesh) and absence (red mesh) of K^+^ show preferential occupancy of the S3 site. E1 maps are contoured at 10σ, E1ATP maps at 11σ, E2P map at 17σ, and E2Pi maps at 12σ. NB: the E2-P state was only imaged in the presence of K^+^.

Enzymatic activity of KdpFABC depends on the presence of the transport substrate K^+^ with a K_M_ of 7 mM for the Q116R mutant (Sweet et al., 2020). We therefore compared density maps from samples in the presence and absence of this ion to look for corresponding conformational changes. However, models created for the various states in the presence of 100 mM K^+^ correlate equally well with density maps for those same states produced in the absence of K^+^ (Fig. 3b, Suppl. Fig. 7). The only observed changes involve specific spherical densities within the SF of KdpA, which is characterized by four canonical sites: S1-S4 (Fig. 3c) (Liu and Lockless, 2013). At the S3 site, we consistently see strong density in maps from samples containing K^+^ (Fig. 3e). This density was also seen in the crystal structure and anomalous scattering was used to confirm assignment as K^+^ (Huang et al., 2017). When K^+^ is absent, the S3 density is much weaker or completely missing in our cryo-EM maps. We also observe density at the S1 site, which was occupied by the guanidino group of Arg116 in the crystal structure. As discussed, the Q116R mutation lowers the apparent K^+^ affinity from micromolar to millimolar, and we previously hypothesized that this effect results from a competition between arginine and K^+^ at this site (Huang et al., 2017). In support of this idea, some of our maps show bifurcated or ambiguous density for the Arg116 sidechain which can be modeled either inserted into the S1 site (e.g., E1), or residing just above it with a K^+^ occupying the S1 site (e.g., E2·Pi with K^+^, Fig. 3d).

## Dimensions of the transmembrane tunnel

The intramembrane tunnel linking KdpA and KdpB is a unique feature of KdpFABC that we previously hypothesized to play a key role in the transport process. To look for changes during the reaction cycle, we plotted radii of the tunnel in the three highest resolution structures (E1, E1·ATP and E2·Pi) starting at the S1 site at entrance to the SF and traversing through this filter, across KdpA and into the Bx site in KdpB (Fig. 4). These plots illustrate the severe constriction of the SF, due to close apposition of carbonyl atoms that serve to distinguish K^+^ from other ions. Once past the SF, the intramembrane tunnel widens into a relatively spacious vestibule that extends about half way to the KdpA-KdpB interface. Comparison of the various structures shows little change in the KdpA region of the tunnel, consistent with the observed rigidity of this subunit. In the E1 structures, the tunnel narrows to ∼1.4 Å as it approaches the subunit interface before widening again in the vicinity of the Bx site. In the E2·Pi structure, movements of M3-M5 in KdpB introduce a bottleneck of ∼0.8 Å and also collapse the cavity next to Bx. Given the sensitivity of this analysis to accurate side chain placements, the lower resolution of KdpB in the E2-P state (4-5 Å) reduces our confidence in the dimension of the tunnel in this region, but we have included the corresponding plot in Fig. 4b (dotted blue line) and Suppl. Fig. 8c for completeness.

**Fig 4.**
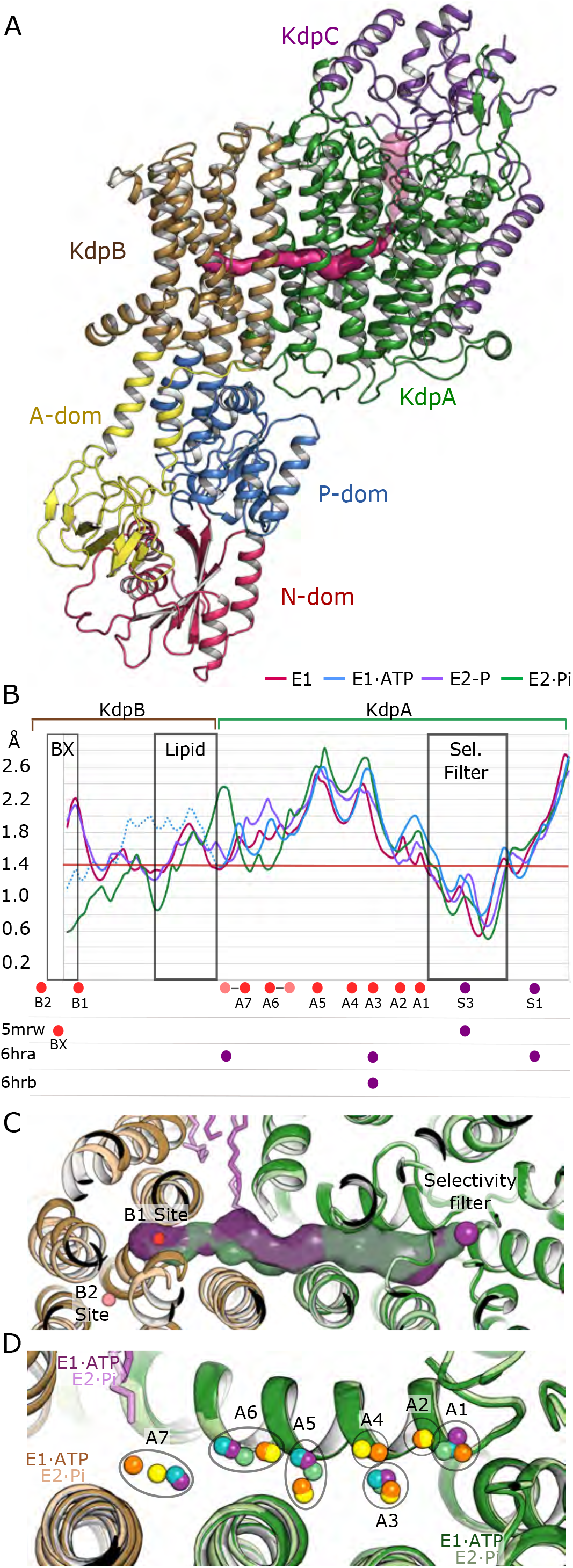
Mapping of the intramembrane tunnel. **(A)** Depiction of the intramembrane tunnel in the E1 state running between the SF in KdpA and the B1 site in KdpB. here in E1 state. Subunit colors are as in previous figures and the tunnel surface is pink. **(B)** Plot of tunnel radii in the four states as indicated in the legend. The E2-P state is dotted because the lower resolution of KdpB makes this analysis less certain. The conventional threshold for water and K^+^ of 1.4 Å is shown as a horizontal red line. The locations of various densities are shown below, both from our density maps (A1-A7, B1, B2, S1, S3) and for previously published structures indicated by their PDBIDs. **(C)** Tunnel surfaces from the higher resolution E1·ATP (purple) and E2·Pi (green) states. The corresponding atomic models are also shown with E1·ATP rendered in darker colors. The purple sticks correspond to a lipid molecule seen in all structures, the purple sphere to K^+^ in the S3 sites and the red sphere to a water in the B1 site. **(D)** Location of densities within the intramembrane tunnel viewed from the periplasm with ribbon models for the E1·ATP (dark colors) and E2Pi (light colors) states. The spheres correspond to densities seen in the higher resolution density maps: E1+K^+^, purple, E1·ATP+K^+^, blue; E2-P+K^+^, green; E2·Pi-K^+^, yellow; E2·Pi+K^+^, orange.

The continuity of the intramembrane tunnel is influenced by a lipid molecule that we modeled at the interface of KdpA and KdpB. The lipid was placed into an elongated density that we observe at this interface in all of our density maps (Suppl. Fig. 8) and that was also observed in previously published cryo-EM maps (Stock et al., 2018). Tentative assignment of lipid is consistent with the bifurcated density visible in our best maps, which was further supported by mass spectrometry of the detergent-solubilized, purified complex used for our cryo-EM experiments (Suppl. Fig. 9). As expected from *E. coli*, phosphatidylethanolamine (PE) and phosphatidylglycerol (PG) with chain lengths of 16 or 18 carbons dominated the mass spectrum. The 16/18 species was most consistent with the densities in the maps (Suppl. Fig. 8b), and we therefore modeled a 16:0/18:1 PE molecule at the interface between M10 of KdpA and M7 of KdpB. This region changes during the E1 to E2 transition, during which M5 from KdpB moves away from the subunit interface and one of the lipid tails is drawn in to fill the resulting space. As a result, the lipid molecule contributes to constriction the tunnel in the E2 states, though it does not affect tunnel radius in the E1 states (Suppl. Fig. 8c). In the absence of this lipid molecule, the intramembrane tunnel would be exposed to the hydrophobic core of the bilayer at the subunit interface, indicating that it plays an important structural role in the complex.

### Character of the intramembrane tunnel of KdpA

The intramembrane tunnel was previously postulated to be either a water-filled cavity for charge transfer or a conduit for K^+^ to move between the subunits (Huang et al., 2017; Stock et al., 2018). We observe several well-defined densities in the tunnels in our density maps allowing us to evaluate these alternative scenarios. Specifically, comparison of our higher resolution maps led to definition of seven distinct sites in KdpA (A1-A7, Fig. 4d, Suppl. Fig. 10) in addition to two new sites in KdpB (B1, B2, Fig. 5). In KdpA, several of the densities are visible in all maps, but as expected, discrete sites are more easily identified in higher resolution maps (Suppl. Fig. 10). Interestingly, the location and spacing of these densities is rather consistent from map to map, regardless of conformational state or presence of K^+^. This result suggests that the tunnel is filled with a loosely ordered array of water molecules throughout the reaction cycle, and the densities have been modeled as such. Near the S4 site of the SF, the tunnel is fairly hydrophilic due to oxygen ligands from side chains of Asn112, Thr230, Ser343, and Asn466. Together with main chain carbonyls from the SF, these side chains have potential to constitute the S4 site described in classical K^+^ channels (Liu and Lockless, 2013). However, S4 is not well formed in KdpA or in its homologs TrkH and KtrB and no density is observed here. The first two densities (A1 and A2) are seen as the tunnel extends away from the SF and widens into a vestibule with radius >2 Å, with side chain oxygens from Ser343, Glu370, Ser378, Tyr381, Thr424 and Asn466 and the main chain carbonyl of Ile421 producing a somewhat hydrophilic environment. This vestibule harbors A3-A6 densities, but lacks any organized coordination geometry that would constitute high-affinity sites. As the tunnel approaches A7, it narrows and becomes distinctly hydrophobic, being lined by side chains from Phe386, Leu389, Ile421, Val538 from KdpA as well as Ala 227 and Leu228 from KdpB. The A7 site is at the subunit interface and thus very near the end of an aliphatic tail from the lipid molecule discussed above. Finally, the tunnel passes hydrophobic side chains on M2 and M5 of KdpB to reach the cavity near Pro264 and the Bx binding site seen in the crystal structure.

**Fig 5.**
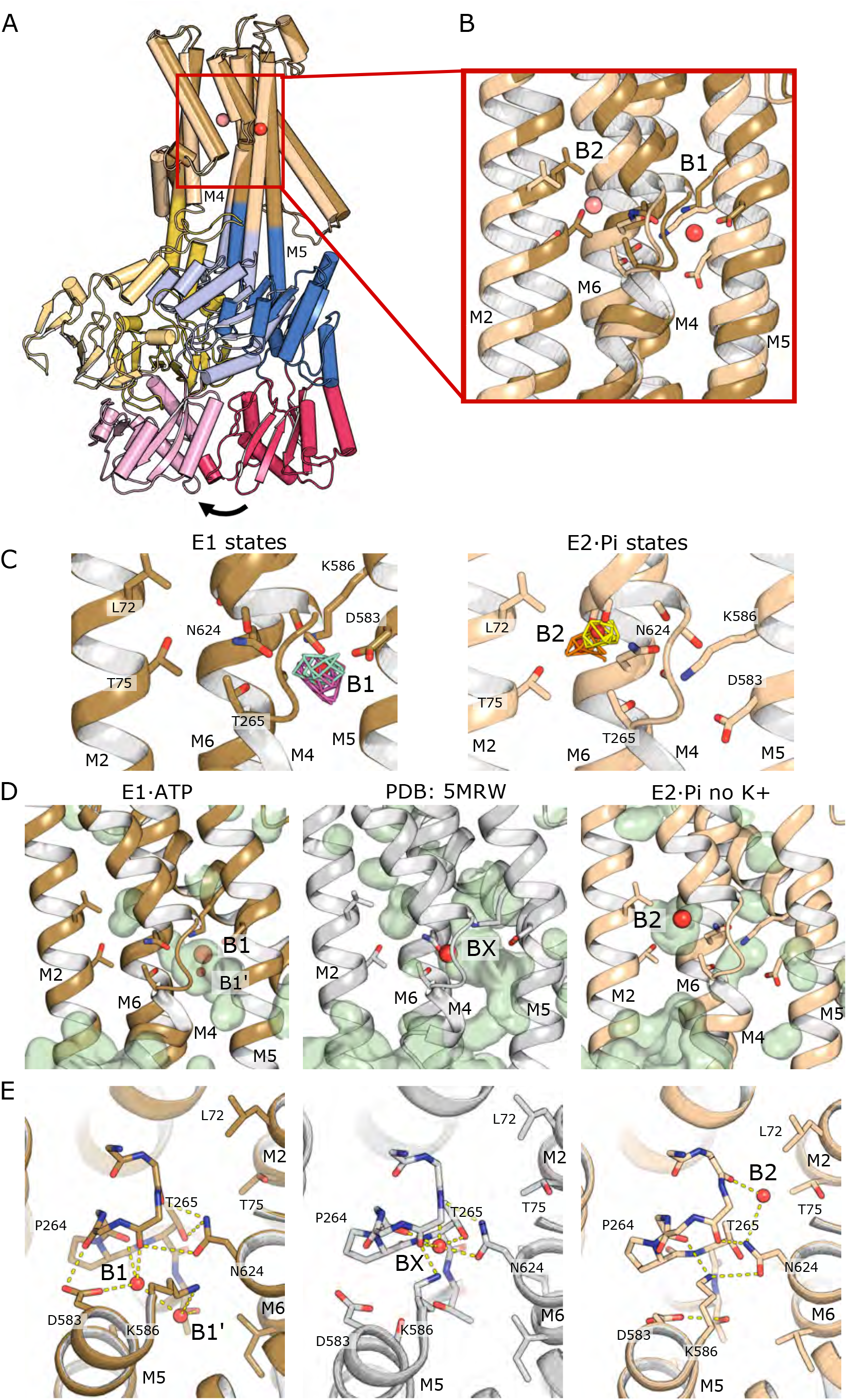
Alternate binding pockets in KdpB. **(A)** Comparison of KdpB conformation in the E1·ATP and E2·Pi states showing large changes in the cytoplasmic domains and movement of M5. This movement of M5 places K586 into the B1 site and thus displaces the ion. The E1·ATP state is rendered in darker colors, which correspond to those in the other figures. Structures were aligned on KdpA and the red box indicates area shown in B. **(B)** Close-up of the B1 and B2 sites showing structural changes associated with the E1 to E2 transition. **(C)** Experimental density in the B1 and B2 binding pockets together with key residues associated with these sites. Map densities are contoured at 6σ and shown as a mesh (E1, purple; E1·ATP, cyan; E2·Pi+K^+^, yellow; E2·Pi-K^+^, orange). The map from the E2-P state is not high enough resolution in this region of KdpB to show distinct densities. **(D)** The cavities in the different states are shown as pale green surface renderings. The solvent radius was set to 1.4 Å for these renderings and water molecules have been modeled at the spherical densities. **(E)** View from the periplasmic side of the membrane showing coordination of water molecules that have been modeled into the spherical densities.

### Spherical Densities in KdpB

In the crystal structure, the Bx site in KdpB is defined by the unwinding of M4 near Pro264 (Fig. 5e) and, here, a water is coordinated by backbone carbonyls of Val260 and Ile263 (M4) and by side chains of Thr265 (M4) and Asn624 (M6). This site is isosteric with the Ca(II) site seen in SERCA (Toyoshima et al., 2000) and the Na(II) site in Na,K-ATPase (Kanai et al., 2013; Nyblom et al., 2013). Our E1 and E1·ATP maps of KdpFABC both display a discrete density in a nearby pocket between M4, M5 and M6, which we have modeled as water in a site we label B1 (Fig. 5). The B1 site is displaced towards M5, relative to Bx, and is close to the Ca(I) and Na(I) sites seen in SERCA and Na,K-ATPase (Suppl. Fig. 11). In our model for E1·ATP, water at the B1 site is coordinated by backbone carbonyls of Cys261 and Ile263 (M4), by the side chain of Asp583 (M5) and by a second water (B1’) that is coordinated by Thr266 (M4), Asn582, Lys586 (M5). This extensive coordination network suggests that the B1 site could have relatively high affinity towards K^+^. Indeed, the K(I) site in Na,K-ATPase is coordinated in a similar way by some of these same residues. Although there is access to the tunnel as described above, the B1 site is otherwise occluded within the membrane domain of KdpB.

The configuration of this region of KdpB undergoes substantial changes in the E2 structures. In E2·Pi, where the resolution is high enough (2.9 Å) for accurate model building, no densities are seen in B1 or its ancillary B1’ site. Indeed, this pocket has collapsed due to a downward movement of M5 that moves the amino group of Lys586 into the B1 site (Fig. 5). Instead, a new spherical density is seen 9 Å away in a site between M2, M4 and M6. The density in this new site, designated B2, has also been modeled as a water that interacts with the backbone carbonyl of Val260 and the amide nitrogen of Asn624; the less extensive coordination suggests that the affinity of this site for K^+^ would be considerably lower than B1. The B2 site is adjacent to a wide, water-filled cavity that opens towards the cytoplasm (Suppl. Fig. 12). This cavity was previously described by Stock et al. and proposed to be an exit site for ions in the E2 state. However, we see this cavity in all of our structures, including the crystal structure, and we were able to identify ordered water molecules in higher-resolution structures of both E1·ATP and E2·Pi states (Suppl. Fig. 12).

## Discussion

In this study, we have used substrate analogs to stabilize KdpFABC in different enzymatic states and have generated the corresponding structures using cryo-EM. Conformational changes seen in KdpB are consistent with four major intermediates in the Post-Albers reaction scheme, but a lack of changes in the other subunits suggest that they remain static during transport. As in previous work, an intramembrane tunnel is observed running from the SF in KdpA to the Bx site in KdpB. For the different reaction intermediates, the architecture of this tunnel is invariant within KdpA, whereas the tunnel undergoes substantial changes in KdpB during the transition from E1·ATP to E2·Pi. Although the tunnel is buried within the protein domains, it has constricted access both to the periplasm through the SF of KdpA and to the cytoplasm via a water-filled cavity in KdpB. Our density maps reveal discrete spherical densities in the SF, in the tunnel and near the Bx site of KdpB; understanding the nature of these densities likely holds the key to understanding the mechanism of transport.

### Assignment of the spherical densities

Although it is not possible to make definitive atomic assignments for densities in a cryo-EM map, the spherical densities in our maps most likely correspond either to water molecules or to K^+^ ions. Comparison of maps derived from structures in the presence and absence of K^+^ provide clues. Within the SF, a strong density is seen at the S3 site in samples containing K^+^ and this density is either absent, or considerably weaker when K^+^ is absent, making assignment at this location straight-forward, while highlighting the ability of both water and K^+^ to bind in these sites (Zhou and MacKinnon, 2003). A second, more variable density is seen in the S1 site, which could be assigned either as K^+^, water, or as the guanidinium group of Arg116 (Fig. 3). Density at the S1 site was also seen in the previous cryo-EM structure of the E1 state (Stock et al., 2018). The construct used in this earlier work carried the wild-type Gln116, supporting the idea that lower affinity produced by the Q116R mutation used here reflects competition for the S1 site ligands at the entrance to the SF. The lower coordination at S1 (5 oxygen ligands) compared to S3 (8 oxygen ligands) suggests that S3 is the main determinant of K^+^ affinity and selectivity, as discussed previously (Huang et al., 2017). Densities have so far not been observed in the S2 or S4 sites, which is consistent with the idea that KdpA is optimized for equilibrium binding of ions, in contrast to the non-equilibrium behavior utilized by channels that promotes rapid exchange between the sites (Liu and Lockless, 2013). In the absence of K^+^, either water or possibly Na^+^ would be expected to bind within the SF to prevent collapse (Jensen et al., 2012). Although Na^+^ binding to the SF is theoretically possible, the strong selectivity of KdpFABC suggests that this ion would not be allowed to pass through the SF and enter into the tunnel.

In contrast to the SF, the map densities within the tunnel itself do not depend on the presence of K^+^, making it likely that these represent water. The region near the SF is hydrophilic and becomes relatively wide (radius ∼2.4 Å) in the vicinity of A3-A6, raising the possibility that a K^+^ ion could move past the water molecules and enter the more hydrophobic and constricted region near the KdpA/KdpB interface (radius ∼1.4 Å). Although there are some polar moieties available to coordinate atoms in the tunnel, they do not constitute an organized binding site, such as seen in the SF. Thus, equilibrium binding by K^+^ in this region is unlikely.

In KdpB, spherical densities are also observed both in the presence and absence of K^+^, but their locations depend on the enzymatic state. In the maps of the E1·ATP state, two densities are observed at the B1 site in the presence of K^+^, but only one density is seen in the absence of K^+^. This observation suggests that the density at the B1 site corresponds to K^+^ and the nearby density (B1’) is a water molecule that helps to coordinate the ion when it is present. Water molecules are similarly seen in SERCA and Na,K-ATPase coordinating their transport substrates (Suppl. Fig. 11). In the E2·Pi state, the B2 site is occupied both in the presence and absence of K^+^. As is discussed below, we believe that B2 corresponds to a low-affinity release site, which could be occupied by K^+^, water or perhaps even Na^+^.

### Comparison with other P-type ATPases

Cryo-EM structures both from the current work and from previous work (Huang et al., 2017; Stock et al., 2018) indicate that KdpA is a static element that serves to select K^+^ from the periplasm and load the transport site of KdpB from within the transmembrane domain. KdpB then functions as an ion pump, like other members of the P-type ATPase superfamily. Thus, comparison of ion binding sites in KdpB, SERCA and Na,K-ATPase sheds light on the mechanism of transport (Suppl. Fig. 11). Both of these other pumps bind multiple cations with a net positive charge of 2-4. As a consequence, the overall neutral charge of the binding site in KdpB is replaced by multiple acidic residues in SERCA and Na,K-ATPase. In particular, Lys586 in KdpB is replaced by a glutamate, and additional aspartate and glutamate residues are present on M4, M6 and M8, all of which participate in ion coordination. From a mechanistic point of view, both SERCA and Na,K-ATPase are designed to release ions to the extracellular side of the membrane in E2-P, whereas KdpFABC releases ions to the cytoplasm. The release in SERCA and Na,K-ATPase is dependent on large movements of M4 and M6 which not only disrupt the coordination network for the ions, but also open an exit pathway. In KdpB, the periplamic sides of the membrane helices do not change much between E1 and E2 states, suggesting that a single K^+^ is simply passed from the B1 to the B2 site where there is a constitutive release pathway present in all of our structures. Despite these conformational differences, there are similarities in the architecture of the ion sites. The B1 site in KdpB, the Ca(I) site in SERCA and the Na(I) site in Na,K-ATPase are all coordinated by homologous residues on M4 and M5 and ancillary water molecules. The Bx site in KdpB is very similar to the Ca(II) site in SERCA and the Na(II) site in Na,K-ATPase, with comparable coordination by main chain carbonyls on M4 and by Asn624 on M6. Asn624 is conserved, Asn796 and Asp800 in SERCA and Na,K-ATPase, and defines a boundary to the ion sites in all three proteins. In SERCA and Na,K-ATPase, the transport ions remain on one side of this boundary, but in KdpB the ion is transferred to the B2 site on the other side of M4. This transfer appears to be the key, energy-dependent step for KdpFABC. The Bx site seen in the crystal structure is half-way between B1 and B2 sites and may therefore be intermediate in this process.

### Structural basis for K^+^ transport

Assignment of densities within the transmembrane domains and conformational changes in KdpB allow us to articulate the steps in the transport cycle (Fig. 6). To begin, ions move through the SF in a constitutive manner with selection of K^+^ over other ions occurring at the S3 site. K^+^ is allowed to transit the tunnel and reach the B1 site in E1 or E1·ATP states, given the uninterrupted nature of the tunnel in these states. In the B1 site of KdpB, K^+^ is coordinated by Asp583 as well as two main-chain carbonyls from M4 and the nearby water molecule in site B1’, which is itself coordinated by Lys586 and Asn582 and Thr266. Asn624 on M6 plays an important role in demarcating this site and holding Lys586 in position. During transition to E2P, K^+^ is transferred to the B2 site on the other side of M4 and Asn624, a site with minimal coordination and presumably lower affinity. Although there is no cavity connecting the B1 and B2 sites in the structures, the Bx site offers an intermediate position that could mediate transfer. The Bx site is occupied in the crystal structure, which is thought to represent an inhibited state, due to phosphorylation of Ser162. Kinetic analyses indicated that phosphorylation of Ser162 trapped the complex in the E1∼P state (Sweet et al., 2020), which is an intermediate between the E1·ATP and E2-P states represented by our cryo-EM structures. Ion binding at the Bx site is therefore likely to mimic an occluded state associated with a true E1∼P intermediate. The positively charged Lys586 plays a key role in the transition to the E2·Pi state as it swings over to occupy the B1 site due to movements in M5; these movements are instigated by the cytoplasmic domains, thereby representing a crucial energy-dependent step in the cycle. The movements also displace Asp583, which is returned to its coordination with Thr266. Both Asp583 and Lys586 were previously identified to be necessary for coupling ATPase activity and K^+^ transport (Bramkamp and Altendorf, 2005). Another consequence of M5 movement is a pinching off of the tunnel at the interface between KdpA and KdpB, thus preventing K^+^ from moving backwards into the tunnel. Once the ion has been delivered to the B2 site in E2-P, it is close to a water-filled pathway leading to the cytoplasm between M1, M2, M3 and M4. Although there is a constriction in our models that would deny direct access of the B2 site to the cytoplasm, it is wider and shorter in the E2·Pi state (Suppl. Fig. 12). The constriction features hydrophilic residues Thr75 and Thr265, which could facilitate escape of the ion, and is lined by M1 and M2 helices. These helices have been shown in other P-type ATPases to move relative to the core membrane helices(Hiraizumi et al., 2019; Sorensen et al., 2004), suggesting that this N-terminal region may have enhanced dynamics, especially because M1 is not tethered to the A-domain in KdpB. It should also be noted that this water-filled exit pathway is present in all observed conformations of KdpB, suggesting that the energy-dependent step is delivery of ions from B1, via Bx, to the low-affinity B2 site, after which they are readily released to the cytoplasm without further changes to the membrane domain.

**Fig 6.**
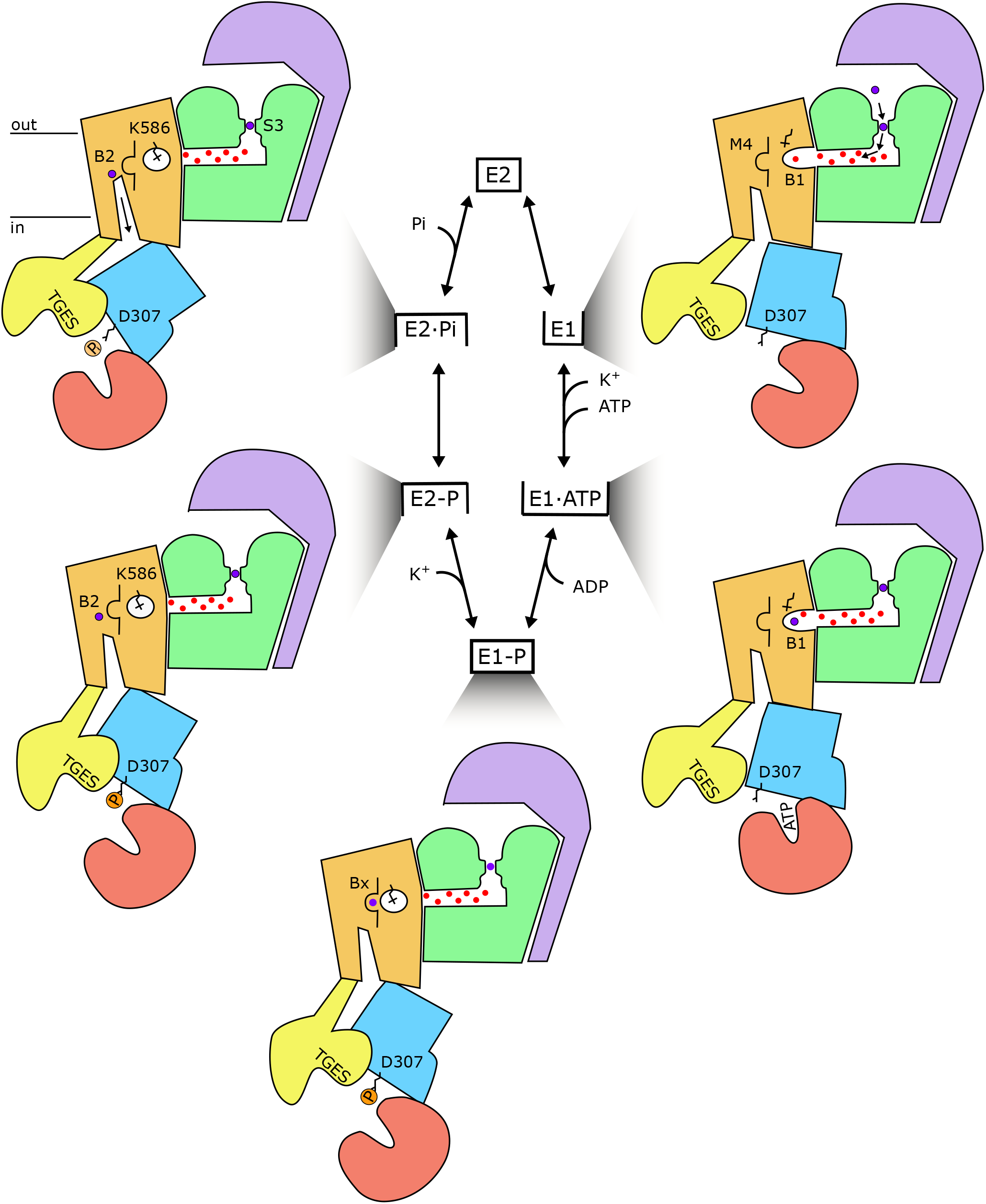
Reaction mechanism of KdpFABC. Cartoon showing functional features and transitions between major intermediates of the Post-Albers reaction cycle. K^+^ enters the SF of KdpA (green) in the E1 state and travels through the water-filled tunnel in the E1·ATP state to reach the B1 site in KdpB (brown). In the E1∼P state, the ion moves to the Bx site, motivated by movement of Lys586 (shown as a crooked stick) and the intramembrane tunnel is pinched off at the subunit interface, thus generating an occluded state. Conformational change accompanying transition to the E2-P state forces the ion into the low-affinity B2 site on the other side of M4 (shown as an open loop). Finally, hydrolysis of the aspartyl phosphate to form the E2·Pi state facilitates release of K^+^ via a pre-formed exit pathway to the cytoplasm.

### Concluding remarks

Cells across all kingdoms of life require high levels of intracellular K^+^, but the transport systems responsible for homeostasis vary. Under normal conditions, K^+^ transport in prokaryotes is handled by SKT and Kup transporters (Epstein, 2003), as is also the case in plants (Dreyer and Uozumi, 2011), while animals use Na,K-ATPase (Clausen et al., 2017). KdpFABC evolved to handle stress conditions and its mechanism is unlike other P-type pumps: it borrows a channel-like subunit from the SKT superfamily to select the transport substrate, which is delivered within the membrane domain to the pump-like subunit from the P-type ATPase superfamily, and then is released into a pre-formed cavity rather than relying on conformational changes to alter the binding site. This partnership appears to be stabilized by accessory subunits, KdpC and KdpF, as well as by a lipid molecule that fills a crucial gap at the subunit interface. Although our mechanistic model defines the basic steps in the reaction cycle, a number of important questions remain. What are the energetics of K^+^ passing through the narrow, hydrophobic part of the tunnel and the transition between B1 and B2 sites? Is the stoichiometry one K^+^ per ATP and does transport of water accompany the transport of K^+^? What are the determinants of the low affinity B2 sites and what are the energetics of release? Finally, what is the allosteric coupling that governs the inhibitory effects of Ser162 phosphorylation in the A-domain. The current work provides a framework to address these questions and to better appreciate the diverse roles of P-type ATPases in biology.

## Acknowledgements

Funding for this work was provided by NIH grant 1R01GM108043 to DLS and by funding from the Independent Research Fund Denmark (grant agreement No. DFF-8021-00161) to BPP.

Electron microscopy was performed at the Cryo-EM Core Facility at NYU Langone Health, with the assistance of William Rice and Bing Wang. In addition, electron microscopy was performed at the Pacific Northwest Center for Cryo-EM at Oregon Health Sciences University, which was supported by NIH grant U24GM129547 and accessed through EMSL (grid.436923.9), a DOE Office of Science User Facility sponsored by the Office of Biological and Environmental Research, and at the Simons Electron Microscopy Center located at the New York Structural Biology Center, supported by grants from the Simons Foundation (SF349247) and NYSTAR.

## Methods

### Cloning and Expression

As in previous work (Huang et al., 2017; Sweet et al., 2020), the kdpFABC operon was cloned into the pBR322 plasmid along with its endogenous promoter and an 8x histidine tag at the C-terminus of kdpC and expressed in E. coli strain TK2499 (F^-^, thi, rha, lacZ, nagA, trkA405, trkD1, Δ(kdpFABC)5, Δ(ompT), from W. Epstein, Univ. of Chicago). We used two different constructs: KdpA-Q116R/KdpB-S162A mutations for cryo-EM work and KdpB-D300A/D302A mutations for functional analysis. For expression of the KdpA-Q116R/KdpB-S162A mutant, a fresh clone of cells were incubated at 37°C overnight in 5 ml of K0 media (46 mM Na_2_PO_4_, 23 mM Na_H_2PO_4_, 25 mM (NH_4_)_2_SO_4_, 0.4 mM MgSO_4_, 6 μM FeSO_4_, 1 mM sodium citrate, 0.2% glucose, 1μg/mL thiamine, 50 μg/mL carbenicillin) supplemented with 5 mM KCl (K5 media). This culture was transferred to 200 mL K1-medium (K0-medium supplemented with 1 mM KCl) and incubated at 37°C for 7 h. The 200 mL cell culture was then transferred to 9 L K0.2-medium (K0-medium supplemented with 0.2 mM KCl) and incubated at 31°C overnight to induce expression of kdpFABC. Cells were harvested at a density of OD_600_= ∼0.8 by centrifugation at 3500g for 20 min at 4°C and cell pellets were either frozen or used immediately for purification. For the KdpB-D300A/D302A mutant, a fresh clone of cells was incubated overnight at 37°C in 10 mL K0-medium. This culture was transferred to 100 mL K0-medium and incubated at 37°C for 4.5 h. When the culture density reached OD_600_ of ∼0.1, and additional 200 mL K0-medium was added and the culture was incubated for another 3 h until OD_600_ returned to ∼0.1. This 300 mL culture was then transferred to 9 L K0-medium and incubated at 31°C overnight. The following morning density had returned to OD_600_ of ∼0.1 and the culture was supplemented with additions of 10 μM KCl every 90 min for 7-8 h until the density reached an OD_600_ of ∼0.4. Cells were then harvested by centrifugation at 3500g for 20 min and the resulting pellet was frozen at -20°C for storage.

### Protein Purification

For purification of KdpFABC, cells were resuspended in 50 mM Tris pH 7.5, 1.2 M NaCl, 10 mM MgCl_2_, 10% glycerol, protease inhibitor tablets (Roche, Basel Switzerland) and 1 mM DTT, and lysed with an Emulsiflex C3 high-pressure homogenizer (Avestin, Ottawa Canada). Whole cells and debris were removed by centrifugation for 15 min at 12,000g, and membranes were pelleted by centrifugation for 2.5 h at 90,140g. Membranes were solubilized by overnight incubation in 50 mM Tris pH 7.5, 600 mM NaCl, 10 mM MgCl_2_, 10% glycerol, 1 mM TCEP and 1.2% n-decyl-β-maltoside (DM) at 4°C using 20 mL per gram of the membrane pellet. After centrifugation for 30 min at 90,140g, the solution was loaded onto a 5 mL Ni-NTA HiTrap chelating column (GE Healthcare, Chicago IL) that was equilibrated in Ni-NTA base buffer (50 mM Tris pH 7.5, 600 mM NaCl, 10 mM MgCl_2_, 10% glycerol, 1 mM TCEP, and 0.15% DM) supplemented with 20 mM imidazole. The column was washed with 20 mL of this same buffer, followed by 20 mL of Ni-NTA base buffer supplemented with 50 mM imidazole. Stepwise 5 mL elutions (collected in 1.5 mL fractions) were then performed as the imidazole concentration was incremented in steps of 50 mM up to a final concentration of 300 mM imidazole. Alternatively, elution was done with a continuous gradient running from 50 - 500 mM imidazole. Fractions containing KdpFABC were pooled, concentrated to ∼0.7 mL using a 100 kDa cut-off Amicon centrifugal filter (Sigma Aldrich, St. Louis MO) and loaded onto a Superdex 200 size exclusion column (GE Healthcare) pre-equilibrated with 100 mM NaCl, 25 mM Tris pH 7.5, 10% glycerol, 1 mM TCEP, and 0.15% DM. Fractions were either used immediately for EM sample preparation, stored for short periods (days) at 4°C for activity assays, or frozen in liquid nitrogen and stored at -80°C.

### Activity Assays

ATPase rates of purified KdpFABC were measured with a coupled enzyme assay (Warren et al., 1974) with 5 μg of protein in a total volume of 0.5 mL at 25°C. The standard assay buffer was composed of 75 mM TES pH 7, 150 mM KCl, 7.5 mM MgCl_2_, 9.6 U/mL lactate dehydrogenase, 9.6 U/mL pyruvate kinase, 2.4 mM ATP, 0.5 mM phosphoenol-pyruvate, 0.36 mM NADH, and 0.15% DM. Concentrations of KCl and ATP were varied for determination of K_M_.

For measurement of K^+^ transport, KdpFABC was reconstituted into proteoliposomes using the method of Levy et al. (Levy et al., 1992). Specifically, a thin film of dry lipid (3:1 weight ratio of E. coli polar extract and 1-palmitoyl-2-oleoyl-sn-glycero-3-phosphocholine (Avanti Polar Lipids, Alabaster AL) was prepared by using a stream of Ar gas to evaporate chloroform from a stock solution (25 mg/ml) followed by 2 h in a vacuum chamber. This dry film was resuspended in transport buffer (20 mM HEPES pH 7.2, 5 mM MgSO_4_, 140 mM K^+^ sulfamate, 1 mM N-methyl-D-glucamine sulfamate) to make a 10 mg/mL lipid stock solution, which was subjected to five cycles of freezing and thawing using liquid N_2_. The stock solution was then extruded 13 times through a 400 nm nucleopore filter (Whatman plc, Maidstone UK) to create homogenous, unilamellar liposomes. Proteoliposomes were made using a lipid:protein weight ratio of 10:1, with 1.25 mg liposomes, 125 μg of KdpFABC, and 375 μg Triton X-100 (Sigma Aldrich) in a total volume of 250 μL. This solution was stirred at room temperature for 30 min followed by addition of 7.5 mg BioBeads (BioRad, Hercules CA) and a further 90 min incubation. Finally, 15 mg BioBeads were added before overnight incubation with stirring at 4°C. Reconstituted proteoliposomes were harvested and bath-sonicated 3 times for 10 sec each before use in the transport assay.

Two different transport assays were used. For the KdpA-Q116R/KdpB-S162A mutant, 25 μL of proteoliposomes were added to a quartz cuvette containing 2 mL of transport buffer. The cuvette was placed in a Fluoromax-4 fluorimeter (Horiba Scientific, Piscataway NJ) at 25 °C with stirring. The voltage-sensing dye DiSC_3_ (Anaspec, Fremont CA) was added (1 μM final concentration from a 1 mM stock in DMSO) and, after a brief equilibration, transport was initiated by addition of 2 mM ATP. Fluorescence was monitored with excitation at 650 nm (5 nm slit) and emission at 675 nm (5 nm slit). Each experiment was terminated by addition of 1.3 μM valinomycin. For the KdpB-D300A/D302A mutant, we used the SURFE^2^RN1 (Nanion Technologies, Livingston NJ). Sensors obtained from the manufacturer were treated with 0.5 mM 1-octadecanethiol in isopropanol and then passivated with 7.5 μg/ml 1,2-diphytanoyl-sn-glycero-3-phosphocholine in n-decane according to the manufacturer’s instructions. Proteoliposomes were diluted 1:10 in non-activating buffer (20 mM HEPES pH 7.2, 300 mM KCl, 5 mM MgCl_2_) and bath sonicated three times for 10 sec each. After adding 50 μL of non-activating buffer to each sensor, 10 μL of the sonicated proteoliposomes were added and the sensors were centrifuged at 2000g for 30 min at room temperature. Measurements of transport involved a 1 sec wash with non-activating buffer followed a 1 sec wash with activating buffer (non-activating buffer supplemented with 1-300 μM ATP) and a final 1 sec wash with non-activating buffer. Quantification of transport was done by plotting peak currents and fitting with the Michaelis-Menton equation using Prism8 software (GraphPad Software, San Diego CA).

### Lipid analysis

An 0.5 mL aliquot containing 1 mg of purified KdpFABC was extracted into chloroform/methanol (Bligh and Dyer, 1959) by dilution in 2 ml methanol followed by addition of 1 ml chloroform and incubation at 40°C for 30 min. Phase separation was then induced by adding additional chloroform and water and by centrifuging the sample at 500 rpm for 5 min. The organic phase was collected and the aqueous phase was re-extracted with chloroform. The combined organic phases were placed in a heating block at 40°C and the solvents were evaporated under a stream of nitrogen. The dry residue was then re-dissolved in 0.2 ml chloroform/methanol (1:1) and analyzed by liquid-chromatography-mass spectrometry (LC-MS/MS) as previously described (Schlame et al., 2020). For this analysis, 7 µl of the extract was injected into a Restek Ultra C18 reversed-phase column (particle size: 3µm, dimensions: 100 × 2.1 mm) and chromatographed at a flow rate of 0.25 ml/min. Solvent A contained 600 ml acetonitrile, 399 ml water, 1 ml formic acid, and 0.631 g ammonium formate. Solvent B contained 900 ml 2-propanol, 99 ml acetonitrile, 1 ml formic acid, and 0.631 g ammonium formate. The chromatographic run time was 30 minutes, with a non-linear gradient of buffer B varying from 30 to 35% (0-2 minutes), from 35 to 67% (2-5 minutes), from 67 to 83% (5-8 minutes), from 83 to 91% (8-11 minutes), from 91 to 95% (11-14 minutes), from 95 to 97% (14-17 minutes), from 97 to 98% (17-20 minutes), from 98 to 100% (20-25 minutes), and from 100 to 30% (25-26 minutes), with the final 4 min at 30%. The column outflow was injected into a QExactive HF-X mass spectrometer (Thermo Fisher Scientific, Waltham MA) operated in negative ion mode. The spray voltage was set to 4 kV and the capillary temperature was set to 350°C. MS1 scans were acquired at a resolution of 120,000, an automatic gain control target of 10^6^, a maximal injection time of 65 ms, and a scan range of 300-2000 m/z. MS2 scans were acquired at a resolution of 30,000, an automatic gain control target of 3×10^6^, a maximal injection time of 75 ms, a loop count of 11, and an isolation window of 1.7 m/z. The normalized collision energy was set to 30 and the dynamic exclusion time to 13 s. MS data were analyzed by the software LipidSearch 4.1 SP1 (Thermo Fisher Scientific, Waltham MA) and the results were curated.

### EM Sample Preparation

Elution fractions from the SEC were used immediately without dilution, except to add inhibitors, K^+^, or Na^+^ as detailed in Suppl. Table 1. The solution of purified protein was generally ∼5 mg/mL and was incubated for 1 hr with one of the following inhibitors: 1mM MgF_4_, 1mM AlF_4_, 2.5 mM BeF_3_, 1 mM VO_4_, 5 mM AMPPCP, after addition of either 100 mM KCl or 100 mM NaCl. After incubation, 3 μL was applied to Ultrafoil grids (R1.2/1.3, Au/Au, 300 mesh, Quantifoil Micro Tools GmbH) that had been glow-discharged for 2 min using a Pelco easiGlow (Ted Pella, Inc, Redding CA). Finally, the grids were blotted and plunge-frozen in liquid ethane using a Vitrobot Mark IV (FEI Corp, Hillsboro OR) at 4°C with a blotting time of 4 sec, blot force of 0, and humidity of 100%.

Stock solutions (10x concentration) of metal fluoride inhibitors were prepared by combining 50 mM NaF with either 10 mM AlCl_3_, 25 mM BeSO_4_, or 10 mM MgCl_2_, followed by incubation at room temperature for 30 min. For the stock solution of orthovanadate, a 200 mM solution of Na_3_VO_4_ was adjusted to pH 10 using NaOH. This solution was incubated at 100°C until the solution became colorless. After cooling to room temperature, this process was repeating until the pH stabilized.

### Image Collection and Processing

Samples were imaged using a Titan Krios G3i microscope (Thermo Fisher Scientific, Waltham MA), running at 300 kV and equipped with a BioQuantum Imaging Filter with either K2 or K3 detectors (Gatan, Inc, Pleasonton CA). Data were acquired with nominal magnifications, calibrated pixel sizes and electron doses listed in Suppl. Table 1 using SerialEM (Mastronarde, 2005), with a defocus range of 1 to 2.2 μm. The resulting datasets were processed with cryoSPARC version 2.9.0 (Punjani et al., 2017) using the following steps. Dose-fractionated movies were normalized by gain reference and corrected for beam-induced motion, using either full-frame followed by local motion correction or patch motion correction only (Rubinstein and Brubaker, 2015). Parameters governing the contrast transfer function (CTF) were estimated using either cryoSPARC’s Patch CTF tool or its implementation of CTFFIND4 (Rohou and Grigorieff, 2015). Templates for particle picking were generated by manually selecting 267 particles followed by 2D classification. Template-based autopicking was then used to locate and extract particles from all micrographs from a given dataset. After 2D classification to remove false positives, 3D density maps were generated through one or two rounds of ab-initio reconstruction, and particles associated with low-population noisy maps were discarded. The remaining particles were further used for heterogenous refinement jobs to generate three classes. Final refinement of promising 3D classes was done using non-uniform refinement jobs, followed by local resolution estimation using cryoSPARC. For the dataset collected in the absence of substrate analogs and K^+^, we applied 3D variability analysis to visualize motions of the cytoplasmic domains. For this analysis, the entire population of 467,590 particles were used to derive three principle components, which were visualized using the “simple mode” with 20 frames per component. Animations comprising the 20 frames were produced using the Volume Series application in Chimera (Pettersen et al., 2004).

### Model Building and Refinement

The crystal structure of KdpFABC (PDBID 5MRW) was used as a starting point for model building. When necessary, individual membrane helices were rigid-body docked and connecting loops were rebuilt using Coot (Emsley et al., 2010). N, P, and A-domains of KdpB were initially docked as rigid bodies for all models. For the E1 and E2P models, the N and A-domains were not altered from rigid-body fits of the crystal structure domains, because the local resolution was not high enough to recognize individual sidechain densities. In other cases, individual residues were adjusted based on the map, except for regions with weak densities. The models were refined in real space using PHENIX (Adams et al., 2010) with the amber force field and iterative cycles of model building and refinement. MolProbity, CaBLAM and Ramachandran Z score (Rama_Z) were used to validate the geometries of the model (Chen et al., 2010; Prisant et al., 2020; Sobolev et al., 2020). Figures displaying the density maps were generated with CHIMERA and those displaying the atomic models were generated with the PyMOL Molecular Graphics System (Schrödinger, LLC).

Although models were build for only five density maps, the conformation of the remaining density maps was evaluated by calculating local cross-correlation coefficients comparing models with maps in comparable conformations. To start, the model was rigid-body docked into the corresponding density map using CHIMERA and then the map-to-model cross-correlation was calculated using the rcc option of the EMDA program included in the CCPEM software suite (Burnley et al., 2017; Wood et al., 2015).

### Analysis of the Internal Cavities

For characterization of internal cavities in KdpFABC, we used Caver Analyst 2.0 Beta (Jurcik et al., 2018). For analysis of the intramembrane tunnel running through KdpB and KdpA, the starting point was defined as the B1 site of KdpB in the E1·ATP model. For each model, we selected the shortest tunnel that ran continuously from the B1 site to S4 site of the SF and then through the SF into the periplasm. For measurements of exit pathway in KdpB, the starting point was set to the B2 site in the E2·Pi model. Settings used for the calculations were as follows. Approximation: 12, Minimum probe radius: 0.5, Clustering threshold: 3.5, Shell depth: 2.5, Shell radius: 4.5, Residues included: AA and 9Y0.

---

**Supp. Fig. 1.**
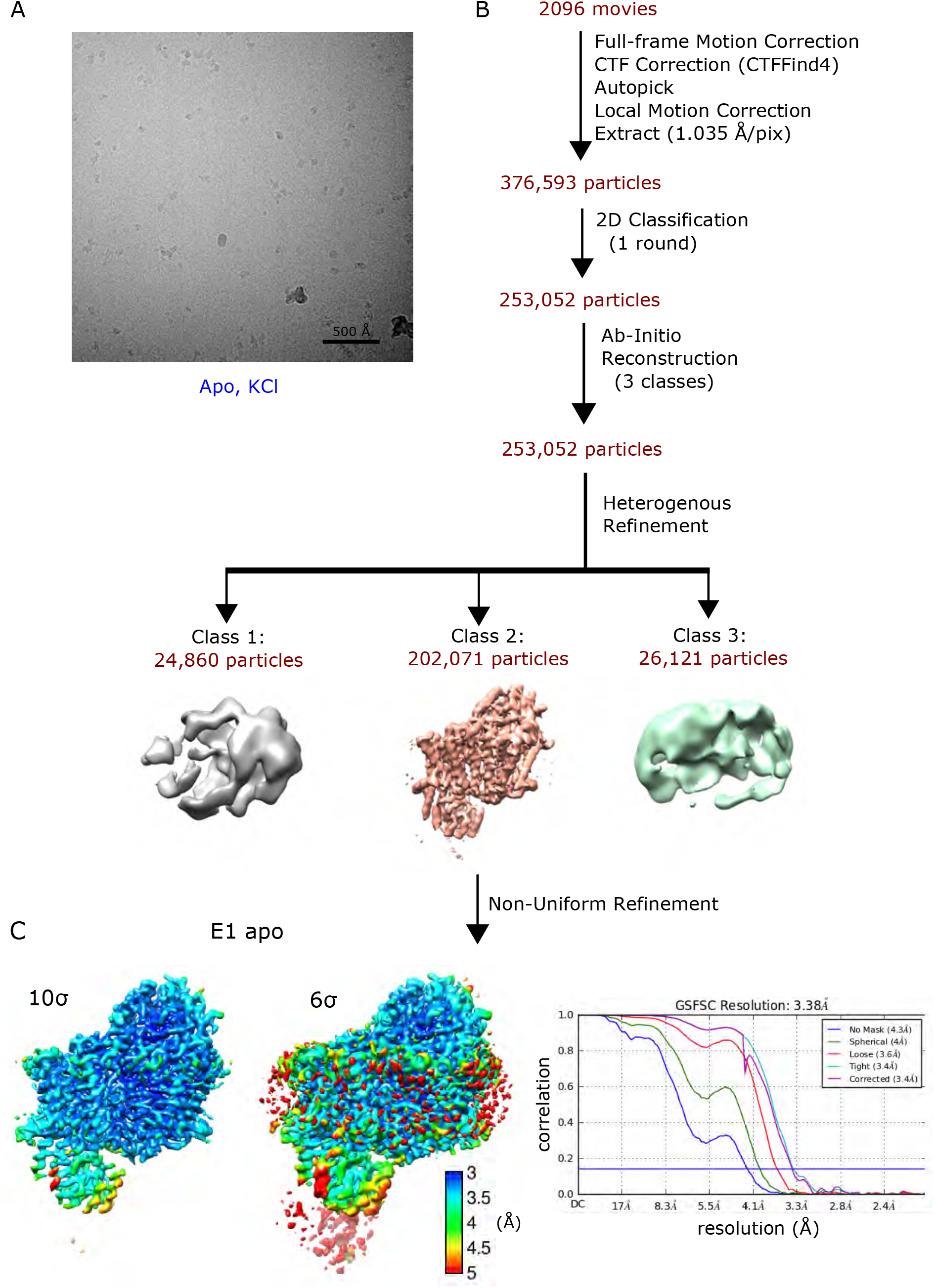
Image processing and reconstruction for samples containing only K^+^. **(A)** Micrograph from Titan Krios microscope using K2 GIF detector. Scale bar corresponds to 500 Å and particle is ∼150 Å in diameter. **(B)** Workflow of image processing and 3D reconstruction in cryoSPARC. **(C)** Sharpened density map from the final non-linear refinement from class 2 colored by local resolution and contoured at thresholds of 10σ and 6σ, in order to see poorly ordered cytoplasmic domains. Corrected curve of the global FSC indicates 3.4 Å based on the 0.14 gold-standard criterion.

**Supp. Fig. 2.**
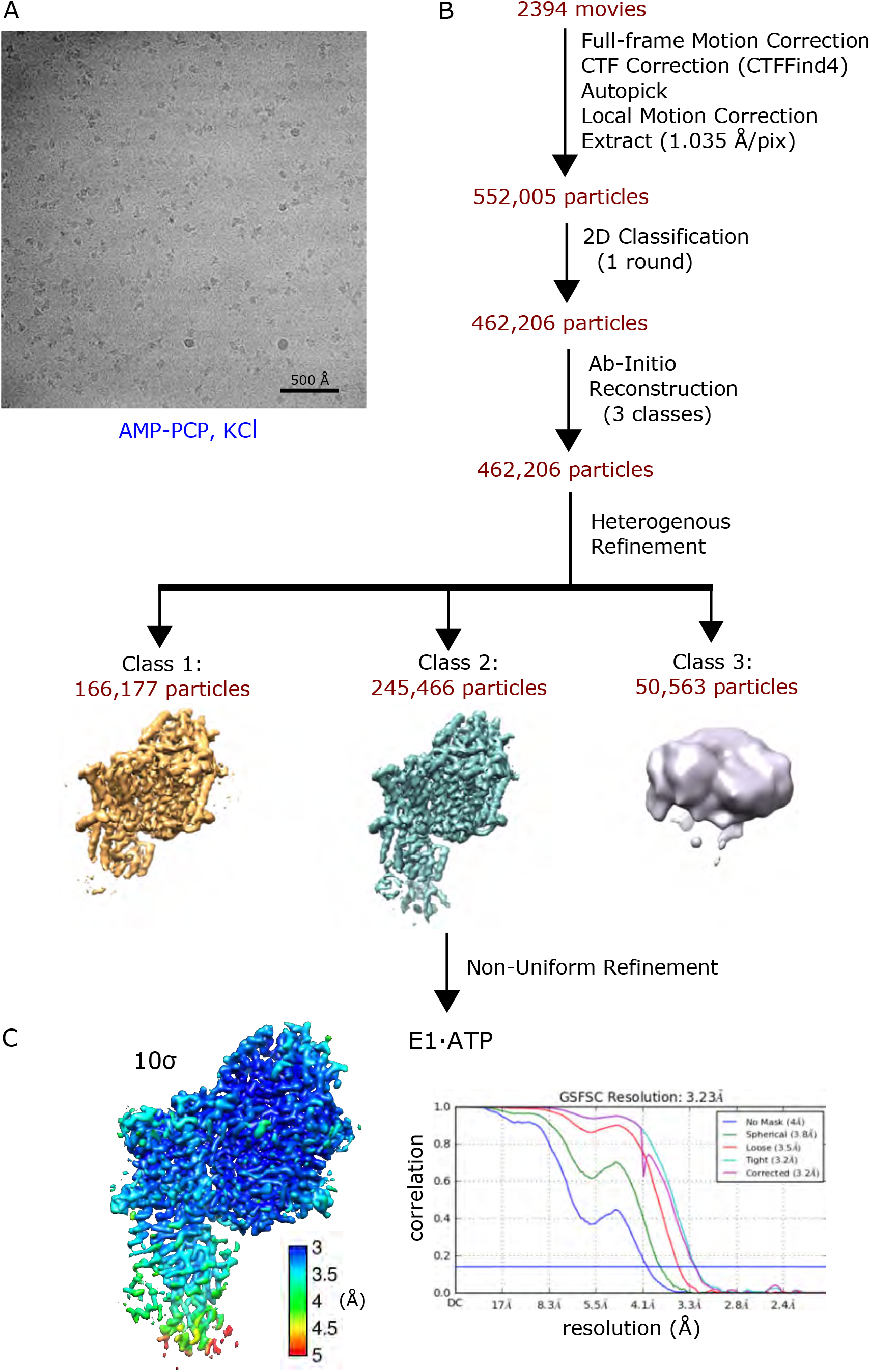
Image processing and reconstruction for samples containing AMP-PCP and K^+^. **(A)** Micrograph from Titan Krios microscope using K2 GIF detector. Scale bar corresponds to 500 Å and particle is ∼150 Å in diameter. **(B)** Workflow of image processing and 3D reconstruction in cryoSPARC. **(C)** Sharpened density map from the final non-linear refinement, contoured at a threshold of 10σ and colored according to local resolution. Corrected curve of the global FSC indicates 3.2 Å based on the 0.14 gold-standard criterion.

**Supp. Fig. 3.**
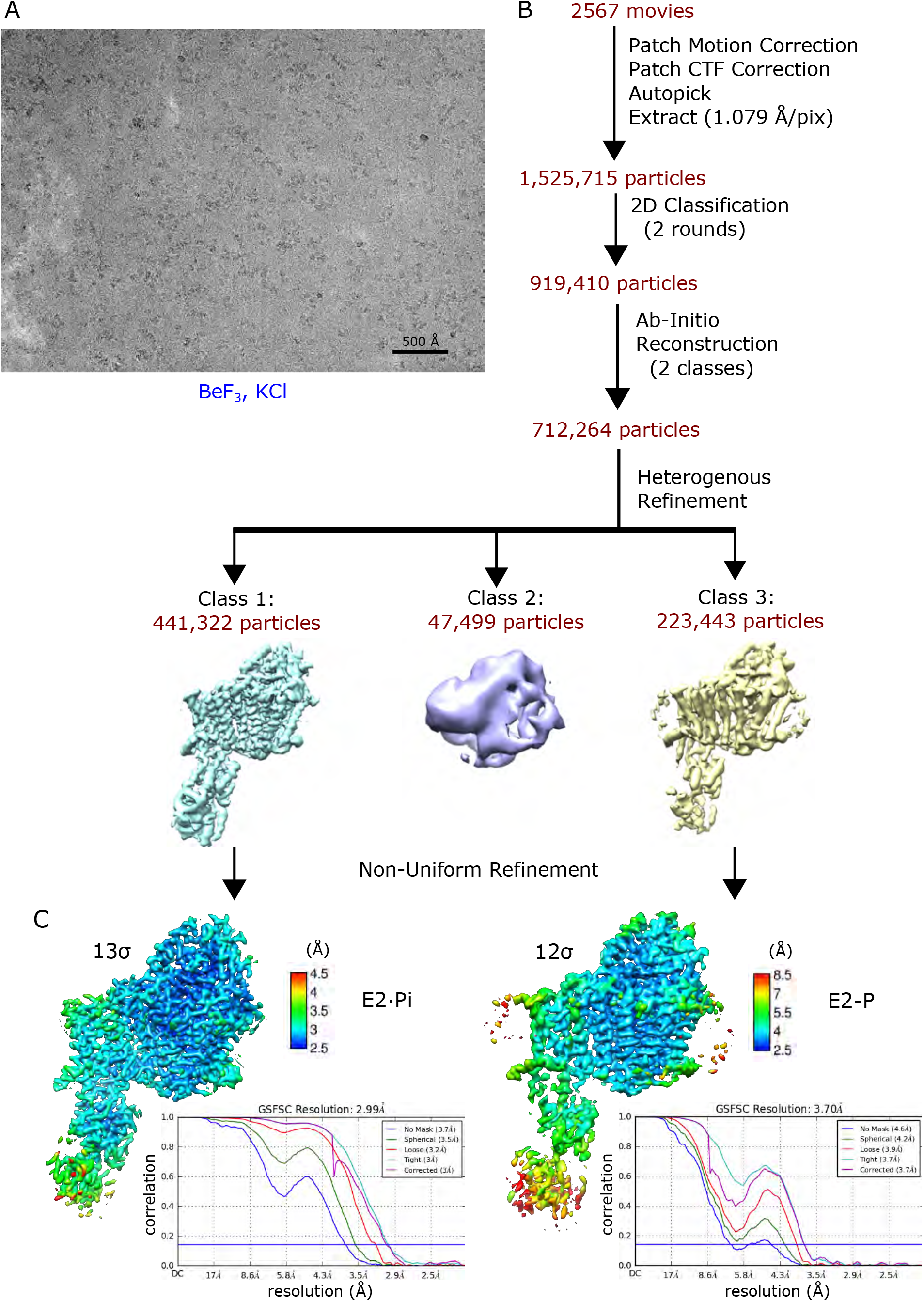
Image processing and reconstruction for samples containing BeF3 and K^+^. **(A)** Micrograph from Titan Krios microscope using K3 GIF detector. Scale bar corresponds to 500 Å and particle is ∼150 Å in diameter. **(B)** Workflow of image processing and 3D reconstruction in cryoSPARC. **(C)** 3D classification produced two distinct structures. Sharpened density maps from non-linear refinements with one corresponding to the E2·Pi state (contoured at 13σ) and to the E2-P state (12σ), colored according to local resolution. Corrected FSC curves indicate global resolutions of 3.0 and 3.7 Å, respectively, based on the 0.14 gold-standard criterion.

**Supp. Fig. 4.**
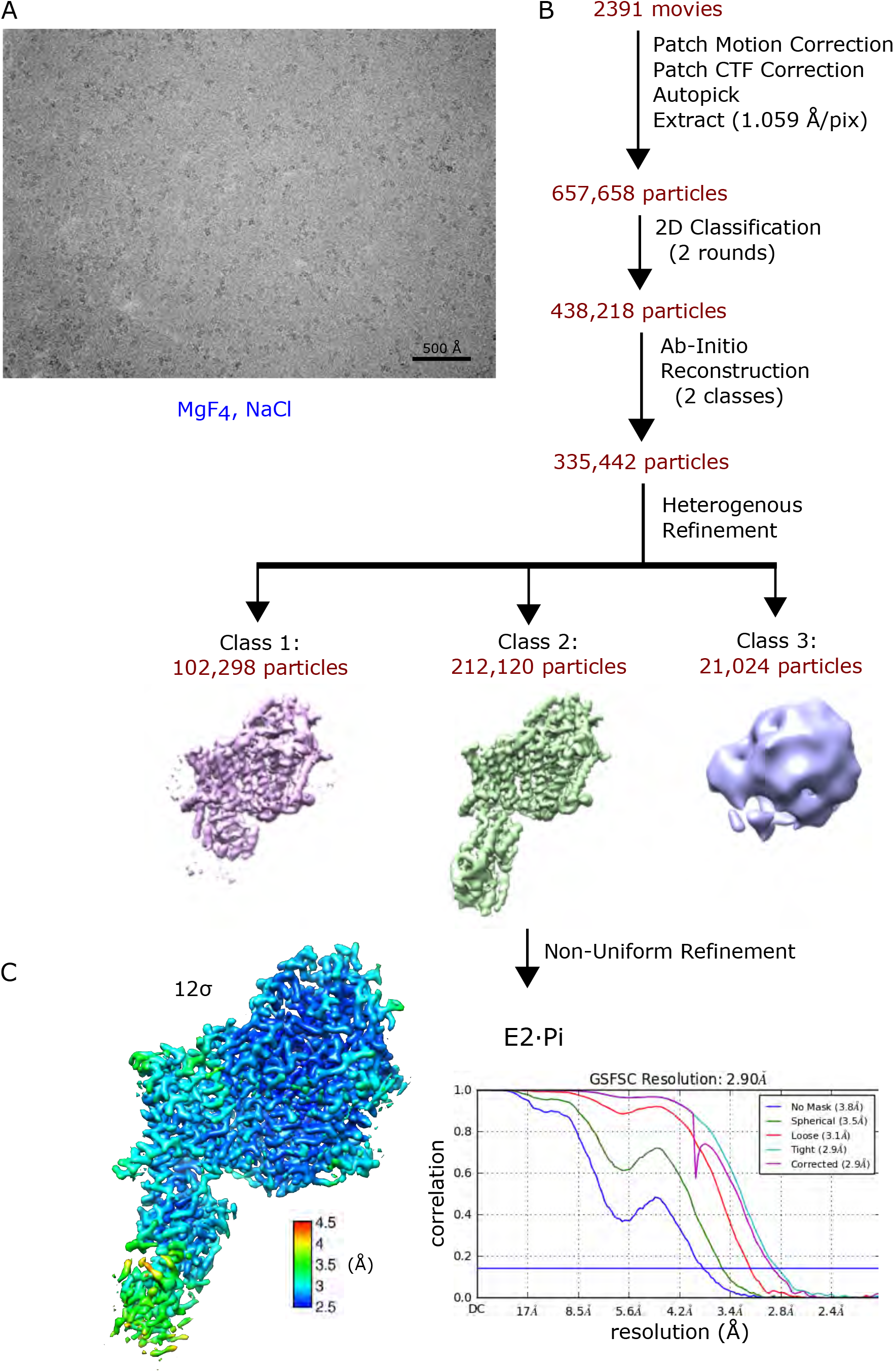
Image processing and reconstruction for samples containing MgF4 without K^+^. **(A)** Micrograph from Titan Krios microscope using K3 GIF detector. Scale bar corresponds to 500 Å and particle is ∼150 Å in diameter. **(B)** Workflow of image processing and 3D reconstrucion in cryoSPARC. **(C)** Sharpened density map from the non-linear refinement colored according to local resolution and contoured at 12σ. Corrected FSC curve indicates a global resolution of 2.9 Å based on 0.14 gold-standard criterion.

**Supp. Fig. 5.**
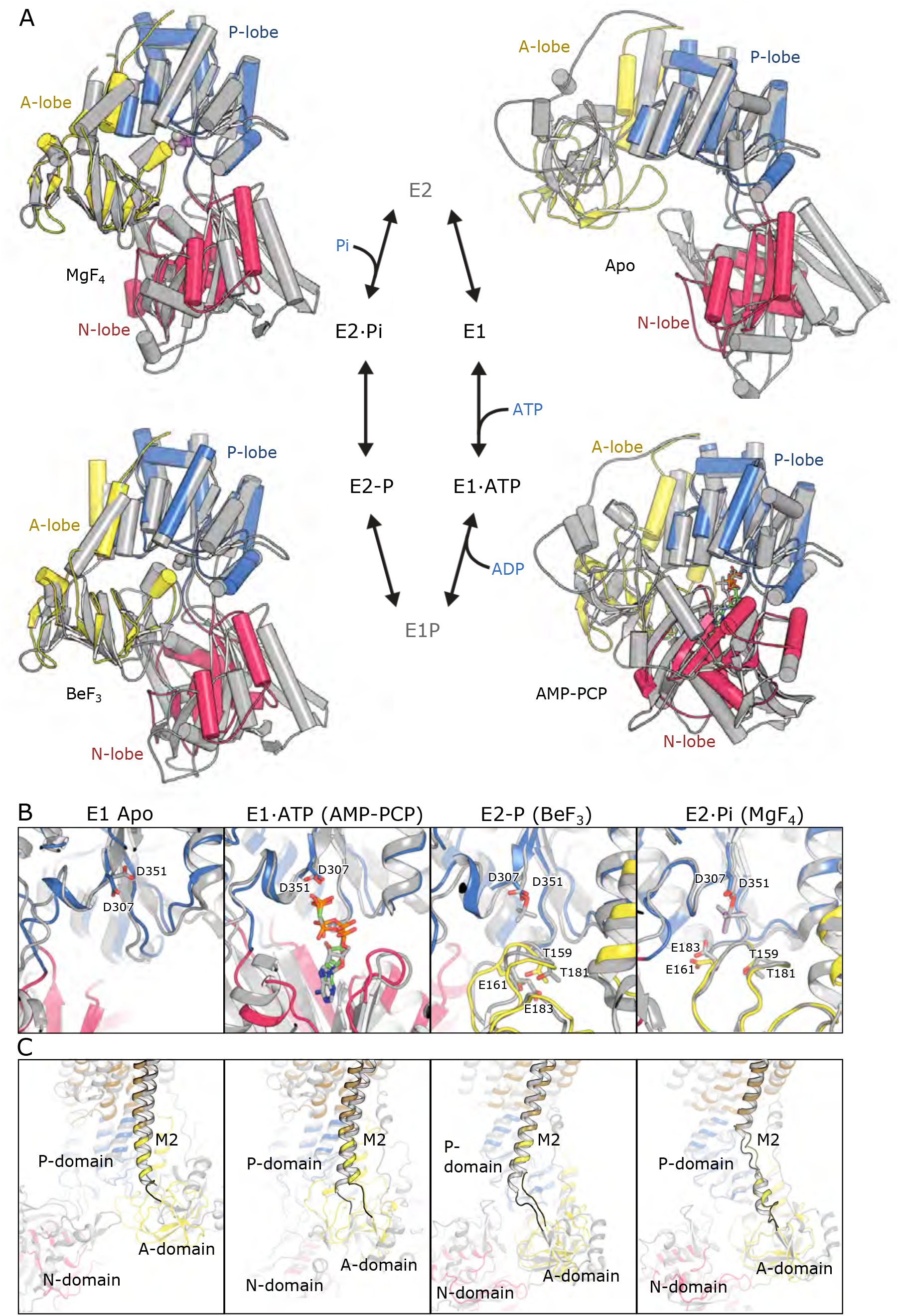
Structural comparison of Kdp and SERCA in matching ligand states. **(A)** Four states are shown which correlate to specific intermediates in the Post-Albers reaction cycle. Cytoplasmic domains of KdpB are colored - N, red; P, blue; A, yellow - whereas those of SERCA are grey (PDBIDs are E1, 1SU4; E1·ATP, 3N8G; E2-P, 3B9B; E2·Pi, 2O9J). Structural alignments are based on the P-domain. AMP-PCP is shown as cyan sticks for KdpB, grey for SERCA. BeF_3_ for SERCA is shown as grey spheres. MgF_4_ is shown as magenta spheres for KdpB and grey for SERCA. **(B)** Closeup of the catalytic sites in the same colors as (A). **(C)** Comparison of the M2 linker in the various states, showing a characteristic unwinding in the E2·Pi state. Domain colors are the same as in (A) with KdpB’s membrane helices colored brown.

**Supp. Fig. 6.**
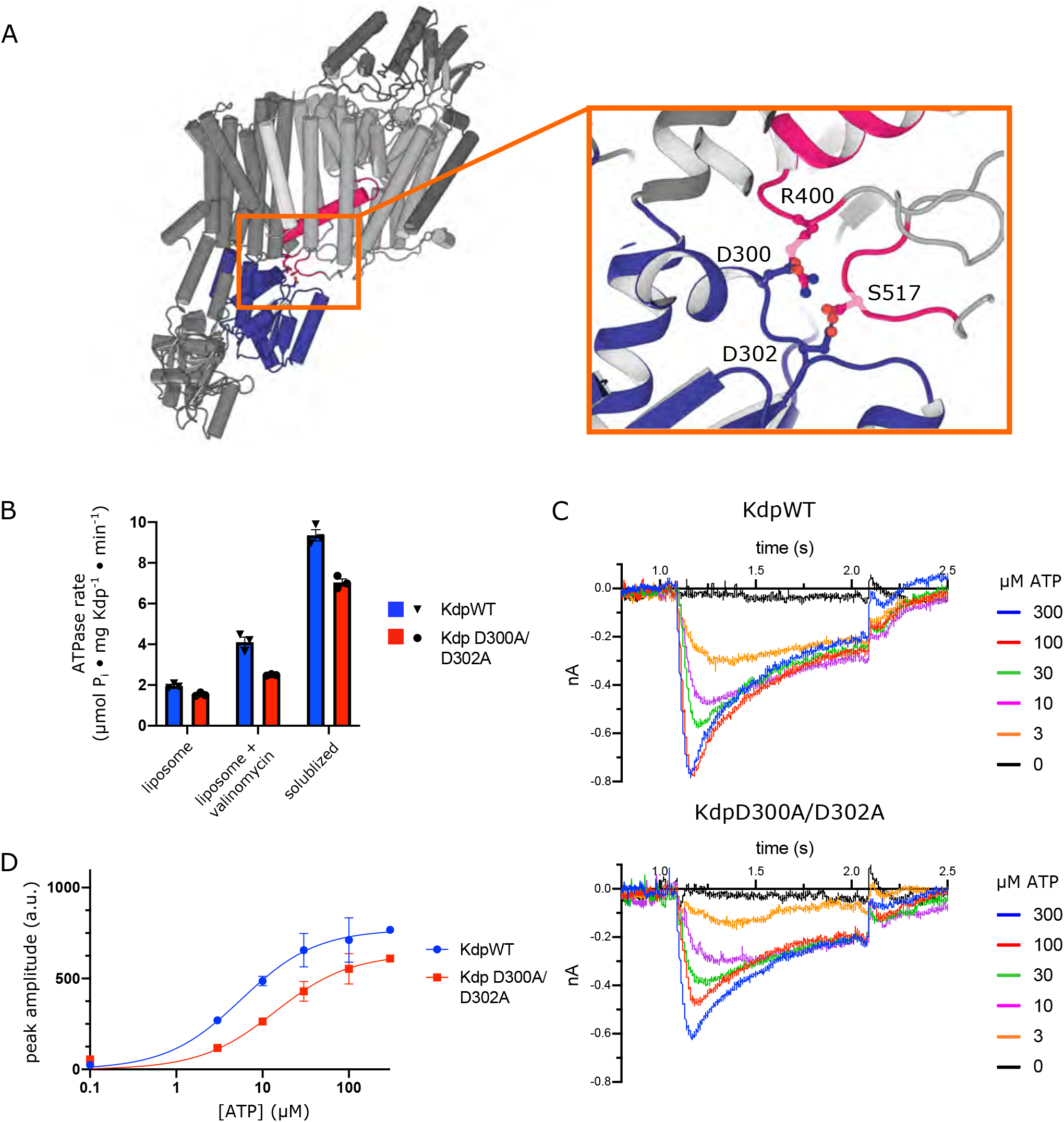
Functional assessment of salt bridge network between KdpA and KdpB. **(A)** Structural overview of the salt-bridge network coupling the P-domain of KdpB (blue) and the helical element D3M_2_ in KdpA (pink), previously postulated (Huang et al., 2017) to relay conformation changes in the former to open a gate for the SF in the latter. **(B)** ATPase assays of purified, preparations of wild-type (KdpWT) and mutant (Kdp D300A/D302A) complexes. The protein was assayed before (solubilized) and after (liposome) reconstitution into proteoliposomes. The latter was assayed in the presence and absence of valinomycin, which stimulates activity by collapsing the membrane potential that otherwise accompanies electrogenic transport. Assuming a 50/50 orientation of KdpFABC in the membrane, two-fold stimulation would be expected for detergent solubilized preparations relative to valinomycin-treated liposomes. **(C)** Raw data for current measured by the SURFE^2^RN1 after initiating transport with ATP at the 1 sec time point. At the 2 sec time point, the sample was washed with ATP-free buffer. **(D)** Aggregated peak currents are fitted with the Michaelin-Menton equation to yield K_M_ of 5.7 mM and 13.9 mM, V_max_ of 770 and 634 sec^-1^ for wild-type and D300A/302A constructs, respectively. Data were measured in triplicate and error bars correspond to S.D.

**Supp. Fig. 7.**
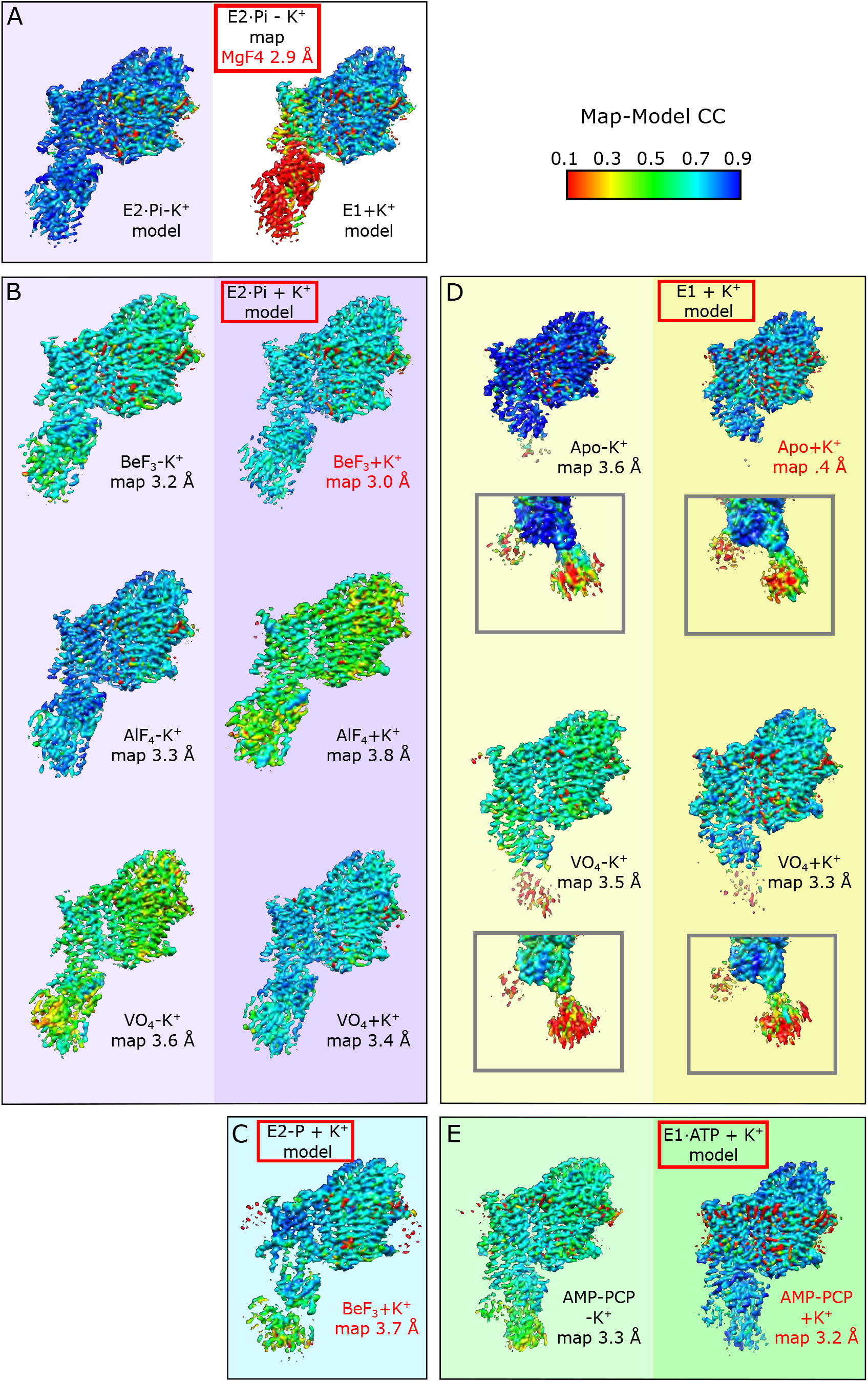
Comparison of structures generated in presence and absence of K^+^. Map-model correlations were used to illustrate addition of K^+^ did not change the observed conformational of KdpFABC. (**A**) The density map for the E2·Pi state in the absence of K^+^ (MgF_4_ without K^+^, see Suppl. Table 1) is compared to atomic models for this same state (left) and for the E1 state with K^+^ (right). Red colors in the cytoplasmic domains as well as yellow/green colors in the membrane domain indicate that the E1 model is a much worse fit compared to the model built for this density map. (**B**) The model built for the E2·Pi state in the presence of K^+^ is compared to six different density maps representing this conformation. These maps were generated from samples with BeF_3_, AlF_4_ and VO_4_ in presence and absence of K^+^. The map used for model building is highlighted in red text. Correlation was dependent on resolution of the maps, but do not suggest conformational changes induced by K^+^. **(C)** There was only one map in the E2-P state, which is compared to its own model. **(D)** The model built for the E1 apo state in the presence of K^+^ is compared with four density maps representing this conformation. The map used for model building is highlighted in red text. Boxed insets show cytoplasmic domains for the corresponding maps at a lower threshold. **(E)** The model built for the E1·ATP state in the presence of K^+^ is compared with two density maps representing this conformation. The map used for model building is highlighted in red text. Correlations were produced with the EMDA program provided with the CCPEM software suite and renderings were made with CHIMERA.

**Supp. Fig. 8.**
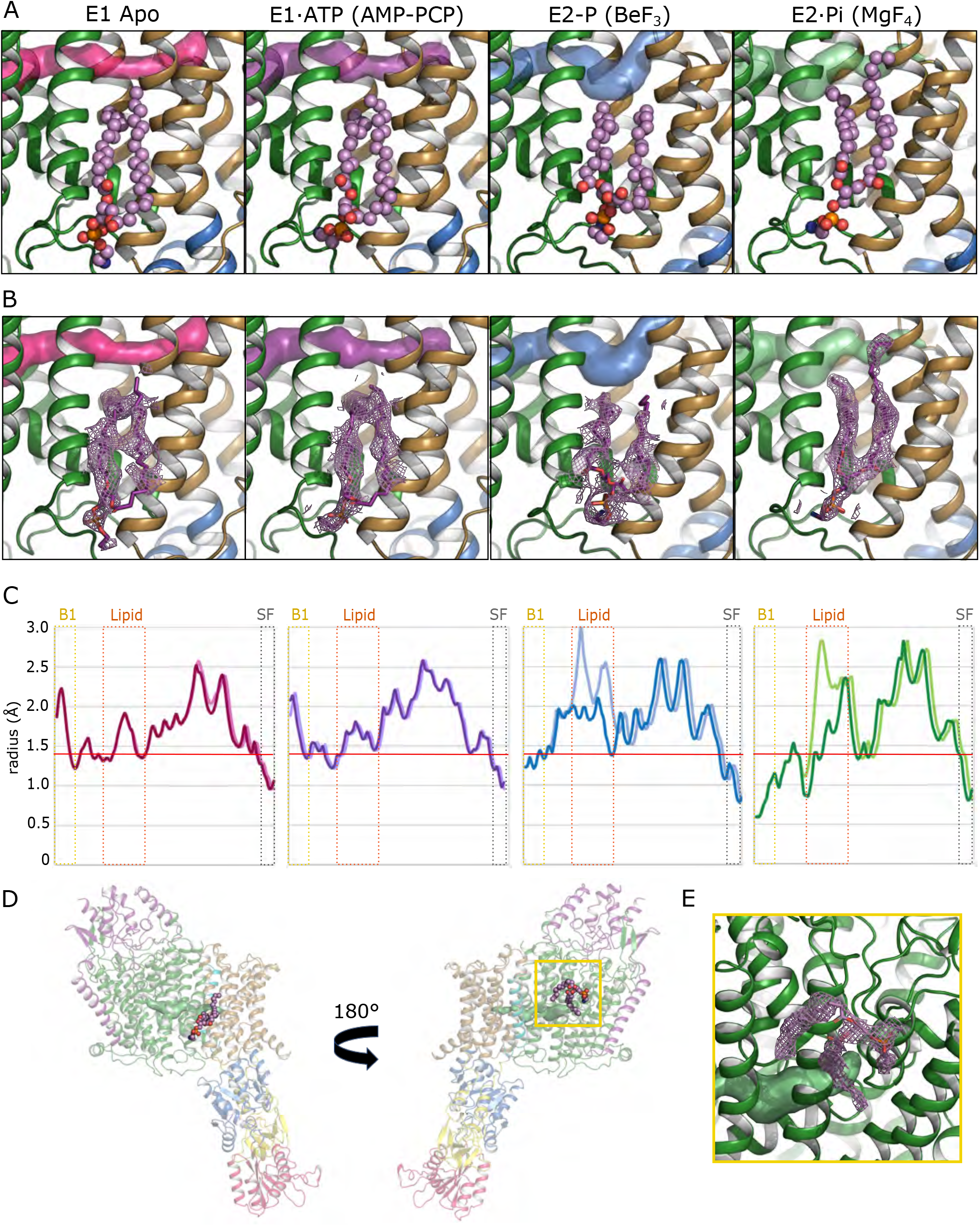
Lipid at KdpA-KdpB interface seals the intramembrane. **(A)** Depiction of a PE lipid molecule (spheres) at the KdpA-KdpB subunit interface in each conformational state. The intramembrane tunnel is shown as a surface at the top of each panel. Secondary structure is colored green for KdpA, brown for the membrane domain of KdpB and blue for the P-domain. **(B)** Map density corresponding to the lipid molecules, which are depicted as sticks. Colors same as (A). Thresholds for the map densities are 7σ for the E1, E1·ATP and E2·Pi states, 7.5σ for E2-P. (C) Tunnel radius plotted for the different states with (darker color) and without (lighter color) the lipid model present in the structure. (D) Overview of KdpFABC showing location of the lipid molecule at the KdpA/KdpB interface (left) and a second PE molecule associated with KdpA on the opposite side of the complex (right). (E) Closeup of the second lipid molecule showing the map density (mesh) as well as the model (sticks). The intramembrane tunnel is also visible as a green surface.

**Supp. Fig. 9.**
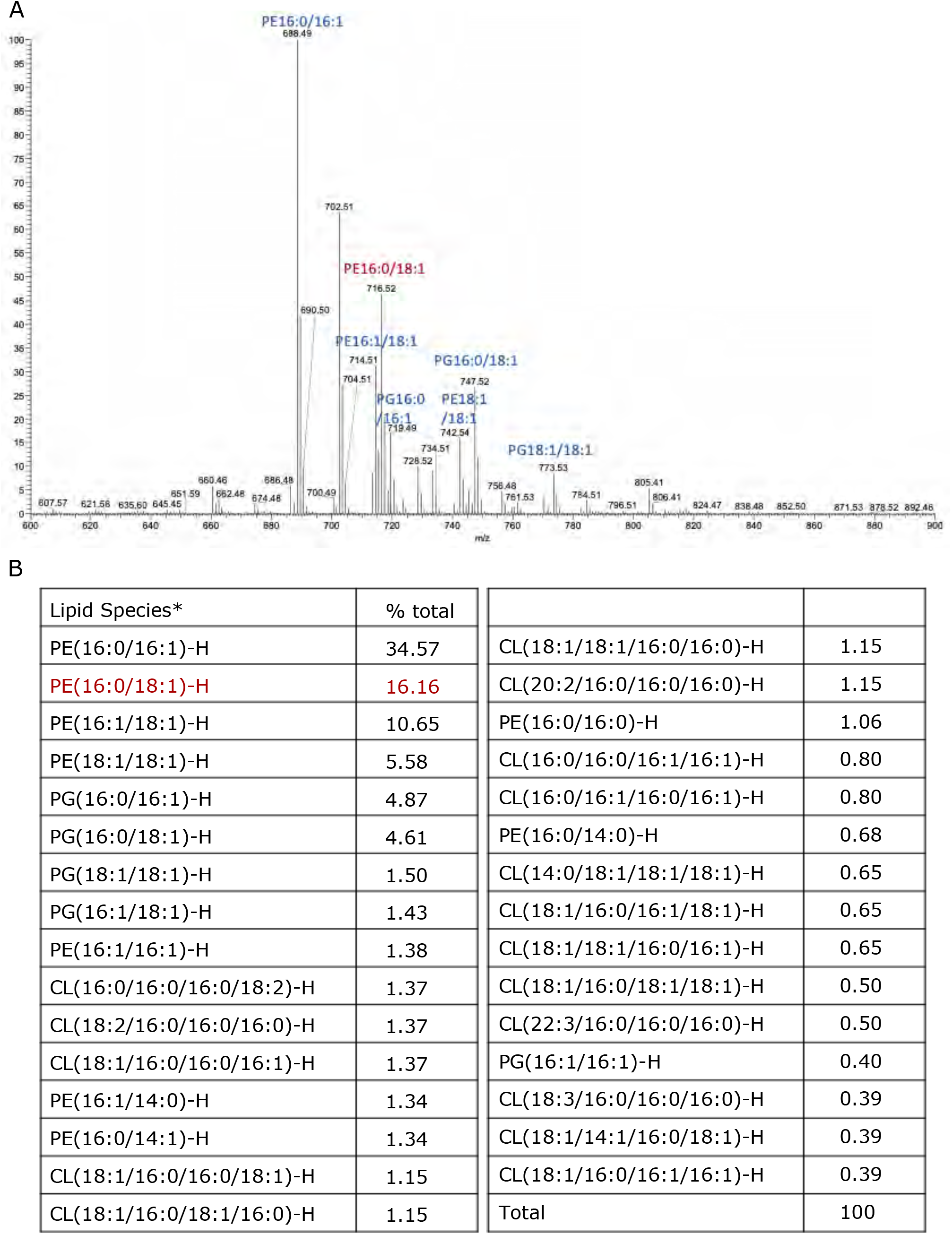
Mass Spectrometry to identify lipid species associated with KdpFABC. **(A)** Spectrum determined by liquid-chromatography-mass spectrometry (LC-MS/MS) of the organic extract from purified KdpFABC complex; the most abundant lipid species are labeled. PE 16:0/18:1 (red label) was consistent with map density at the KdpA/KdpB interface. **(B)** Listing of lipid species identified by LC-MS/MS and quantified according to their MS1 intensity. Lipids are specified by the shorthand notation (Liebisch et al., 2013), in which x:y denotes a fatty acid with × carbon atoms and y double bonds. CL, cardiolipin; PE, phosphatidylethanolamine; PG, phosphatidylglycerol. Abundance is given as percent of total lipid ion current.

**Supp. Fig. 10.**
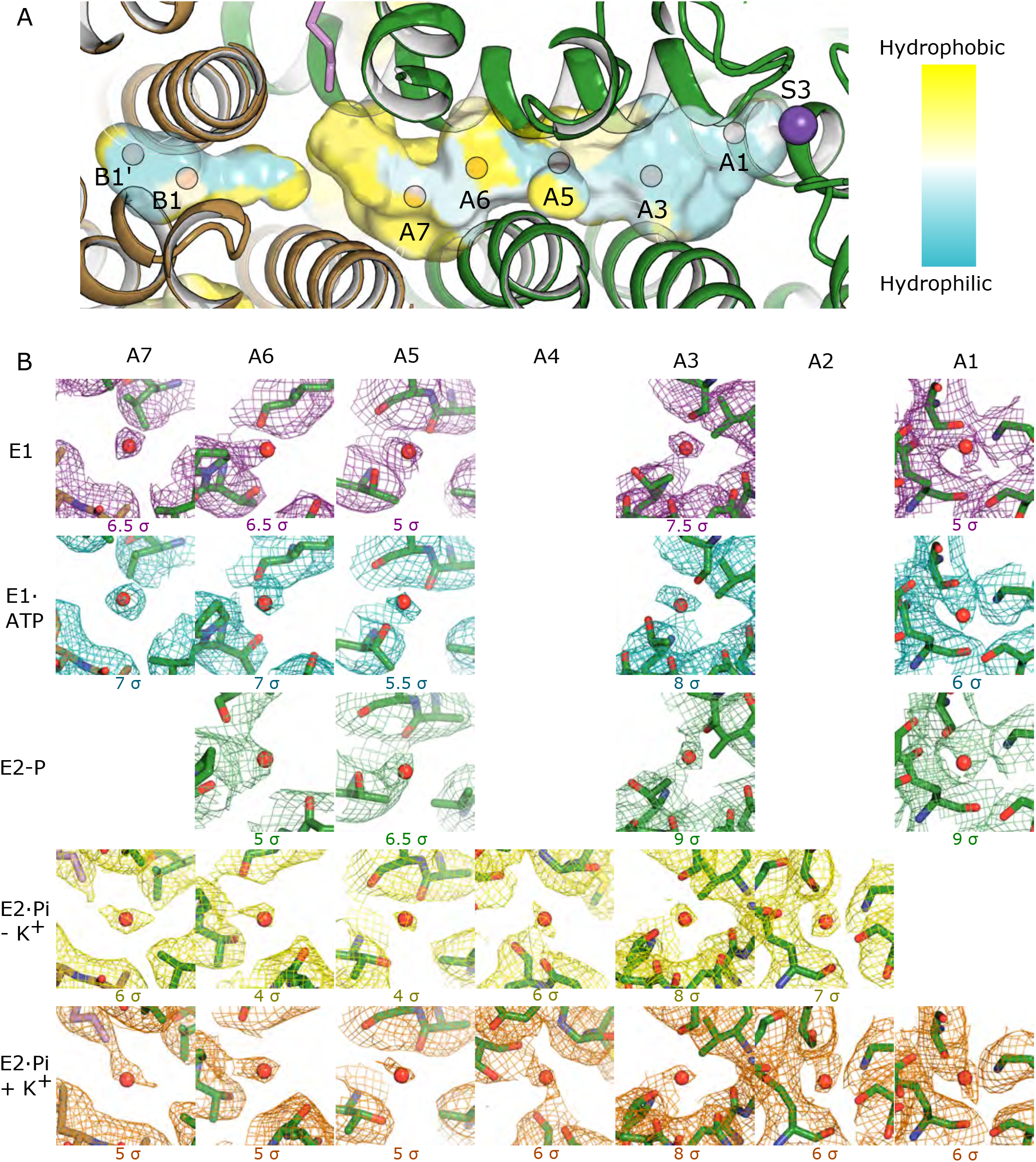
Water molecules in the intramembrane tunnel. **(A)** Surface of the intramembrane tunnel colored according to residue hydrophobicity ranging from cyan to yellow as shown in the key. Location of sites as discussed in the text. **(B)** Map densities associated with water molecules in the various structures. Each mesh was generated with the indicated threshold values. A missing panel indicates a lack of density in the corresponding density map. Mesh colors are consistent with spheres in Fig. 4d.

**Supp. Fig. 11.**
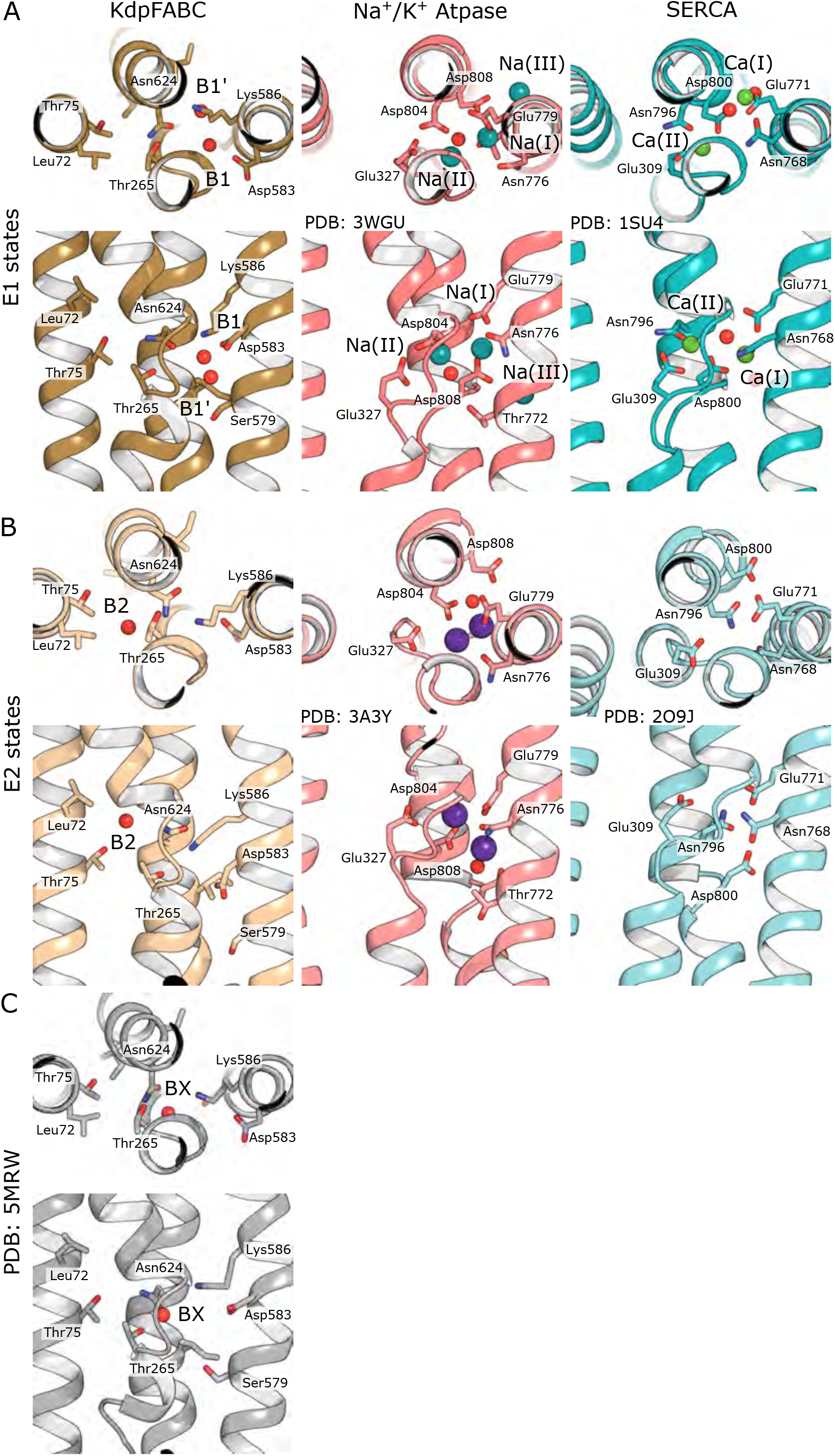
Comparison of ion binding sites from KdpFABC, Na,K-ATPase and SERCA. **(A)** Ion binding sites in E1 conformations viewed from the extracellular side of the membrane (top) and parallel to the membrane plane (bottom). The KdpFABC structure corresponds to the E1·ATP state. The Na,K-ATPase structure (PDBID 3WGU) is in the E1∼P transition state stabilized by ADP and AlF_3_. The SERCA structure (PDBID 1SU4) was solved in the presence of Ca^2+^. (**B**) Ion binding sites in E2 conformations viewed from the extracellular side of the membrane (top) and parallel to the membrane plane (bottom) The KdpFABC structure corresponds to the E2·Pi state in the absence of K^+^. The Na,K-ATPase structure (PDBID 3A3Y) is stabilized by ouabain and K^+^. The SERCA structure (PDBID 2O9J) is stabilized by MgF_4_ and cyclopiazonic acid. (C) Ion binding site seen in the crystal structure (PDBID: 5MRW) in the same orientations as the other panels.

**Supp. Fig. 12.**
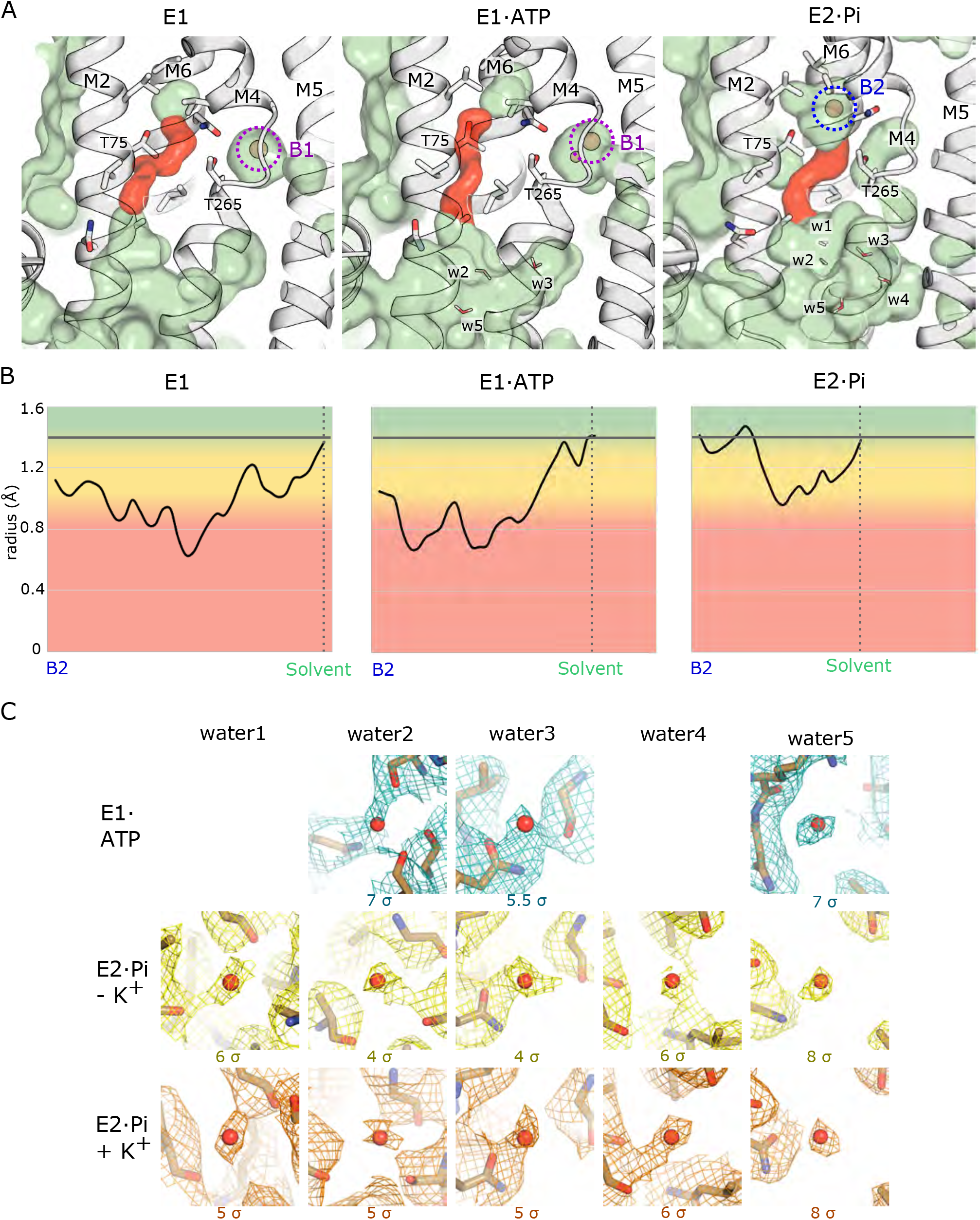
Exit path from the B2 site to the cytoplasm. **(A)** Views of B1 and B2 sites in three different states. Membrane helices of KdpB are shown in white. Cavities are shown in pale green, except for the constricted pathway (red) that connects the B2 site to a wider, water-filled cavity leading to the cytoplasm. Thr75 and Thr266 line the constriction, potentially facilitating escape of an ion bound at B2. Water molecules within this cavity are labeled as w1-w5. (**B**) Radius of the constricted pathway connecting the B2 site to the water-filled cavity (red in A). Shading indicates feasibility of an ion escaping through this constricted region, with the conventional cutoff of 1.4 Å indicated by a horizontal line. (**C**) Map densities associated with water molecules seen along the cytoplasmic exit path at the indicated thresholds.

**Suppl. Table 1.**
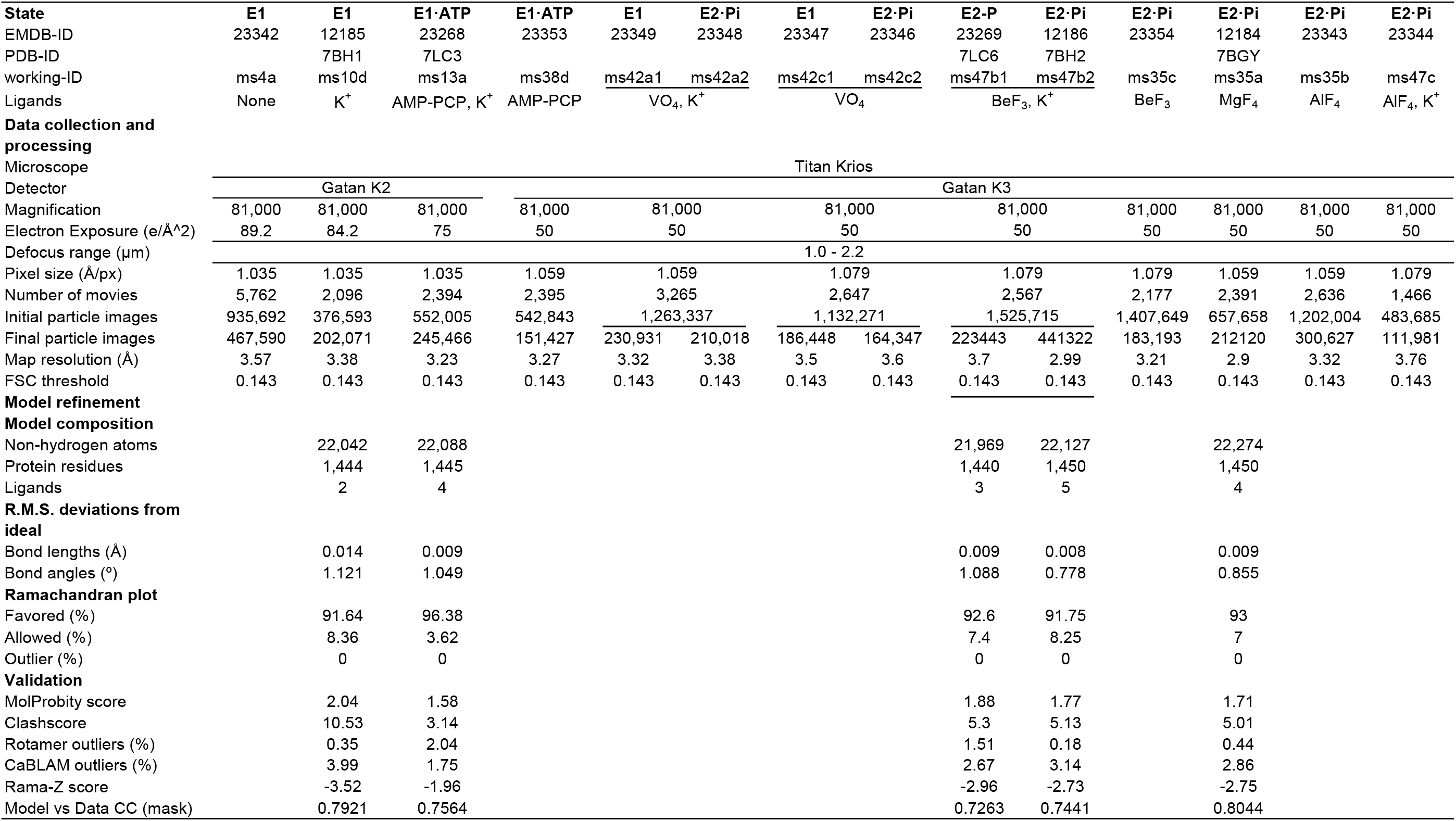
Statistics for collection and processing of all cryo-EM datasets Comparison of structures generated in presence

**Suppl. Movie 1.**

Animation derived from 3D variability analysis of the data set produced in the absence of substrate analogs and K^+^. This animation comprises 20 structures derived from component 0 and illustrates the innate mobility of cytoplasmic domains in the absence of ligands. A low threshold has been used in order to show the cytoplasmic domains; thus, the membrane domains are shrouded in an ellipsoidal micelle.

**Suppl. Movie 2.**

Animation derived from 3D variability analysis of the data set produced in the absence of substrate analogs and K^+^. This animation comprises 20 structures derived from component 1 and illustrates the innate mobility of cytoplasmic domains in the absence of ligands. A low threshold has been used in order to show the cytoplasmic domains; thus, the membrane domains are shrouded in an ellipsoidal micelle.

**Suppl. Movie 3.**

Animation derived from 3D variability analysis of the data set produced in the absence of substrate analogs and K^+^. This animation comprises 20 structures derived from component 2 and illustrates the innate mobility of cytoplasmic domains in the absence of ligands. A low threshold has been used in order to show the cytoplasmic domains; thus, the membrane domains are shrouded in an ellipsoidal micelle.

